# Protein interaction kinetics delimit the performance of phosphorylation-driven protein switches

**DOI:** 10.1101/2023.11.06.565761

**Authors:** Daniel L. Winter, Adelgisa R. Wairara, Jack L. Bennett, William A. Donald, Dominic J. Glover

## Abstract

Post-translational modifications (PTMs) such as phosphorylation and dephosphorylation can rapidly alter protein surface chemistry and structural conformation which can, in turn, switch protein-protein interactions (PPIs) within signaling networks. Recently, *de novo* designed phosphorylation-responsive protein switches have been created that harness kinase- and phosphatase-mediated phosphorylation that modulate PPIs. PTM-driven protein switches could be useful for investigating PTM dynamics in living cells, developing biocompatible nanodevices, and engineering signaling pathways to program cell behavior. However, little is known about the physical and kinetic constraints of PTM-driven protein switches, which limits their practical application. In this study, we present a theoretical framework to evaluate two-component PTM-driven protein switches based on four performance metrics: effective concentration, dynamic range, response time, and reversibility. Our computational models reveal an intricate relationship between the binding kinetics, phosphorylation kinetics, and switch concentration that governs the sensitivity and reversibility of PTM-driven protein switches. Building upon the insights of our theoretical investigation, we built and evaluated two novel phosphorylation-driven protein switches consisting of phosphorylation-sensitive coiled coils as sensor domains fused to fluorescent proteins as actuator domains. By modulating the phosphorylation state of the switches with a specific protein kinase and phosphatase, we demonstrate fast, reversible transitions between easily differentiated “on” and “off” states. The response of the switches linearly correlated to the concentration of the kinase, demonstrating its potential as a biosensor for kinase measurements in real time. As intended, both switches responded to specific kinase activity with an increase in fluorescence signal and our model could be used to distinguish between two mechanisms of switch activation: dimerization or a structural rearrangement. In summary, the protein switch kinetics model presented here should be useful to guide the design of PTM-driven switches and tune their performance towards concrete applications.

## Introduction

The modulation of protein-protein interactions (PPIs) by post-translational modifications (PTMs) underpins cell signaling pathways that mediate appropriate response to intracellular and extracellular stimuli. Cells use PTMs such as protein phosphorylation to integrate multiple signaling pathways and regulate cellular responses, such as cell division. This is achieved by employing protein-modifying enzymes to rapidly alter the surface chemistry of proteins via the addition and removal of chemical moieties, for example, through the action of protein kinases and phosphatases that add or remove phosphate groups, respectively. The addition of PTMs such as phosphorylation can alter the charge and steric bulk of individual residues, either of which can affect protein function through changes in protein structure or PPIs. Ultimately, PTM-mediated protein signaling is an interplay between the kinetics of PTM addition and removal and the kinetics of PTM-driven structural rearrangements, including the modulation of PPIs^1,2^.

Recently, the central function of PTMs as a molecular signaling (or “information-processing”) mechanism has been harnessed for the *de novo* design of protein switches that dynamically respond to phosphorylation by specific kinases and, in some cases, to dephosphorylation by phosphatases^3–6^. Protein switches that are modulated by enzymatic inputs are promising tools to enable the investigation of PTM dynamics in cells^7,8^, the development of biocompatible nanodevices that sense PTM biomarkers for diagnosis or treatment of diseases, and the engineering of orthogonal signaling pathways to program the behavior of synthetic organisms^9^.

We have previously described the design of interacting peptides whose binding affinities could be modulated by the activities of protein kinase A (PKA) and lambda protein phosphatase (λPP)^3^. Two of the designed peptide pairs assembled as heterodimeric coiled coils whose interactions were strengthened following phosphorylation by PKA. This increase in affinity was also reversible through dephosphorylation by λPP. Other research groups have subsequently reported the creation of protein switches based on similar design strategies, but where phosphorylation disrupted either coiled coil interactions^4,6^ or the conformation of an α-helix bundle based on the LOCKR protein switch system^5^. Prior to the recent efforts focused on *de novo* designed protein switches, naturally occurring PTM-binding domains such as the 14-3-3 domain and the forkhead-associated 1 (FHA1) domain have been exploited to build protein devices that can produce a fluorescent signal in response to kinase activity^10–12^.

Despite the potential of PTM-driven protein switches for reprograming cell behavior, a theoretical framework to guide the design of protein switches – and, more broadly, to investigate the kinetics of PTMs and their dynamic effects on PPIs – is lacking. Phosphorylation-driven protein switches have been designed through semi-rational approaches and without specific performance metrics such as the dynamic range, effective concentration, response time, and reversibility of the protein switches (Figure 1). Fundamentally, a two-component protein switch functions by changing the affinity between its component subunits, and thereby the proportion of subunits that are interacting. However, at too high or low concentration, the subunits will be assembled or disassembled regardless of the stimuli. Thus, to fully realize the potential of PTM-driven switches and their many applications, these metrics should be considered in the design process to match to the desired switch application. The importance of explicitly recording performance metrics of a protein switch has been recognized in the systematic screening of PQQ-glucose dehydrogenase (GDH)-based protein switches to detect calmodulin-binding peptides^13^. The authors investigated three chosen metrics (dynamic range, response time, and reaction rate) of over 200 variants of the protein switch and found that trade-offs existed between these metrics for the GDH-based switch.

**Figure 1.**
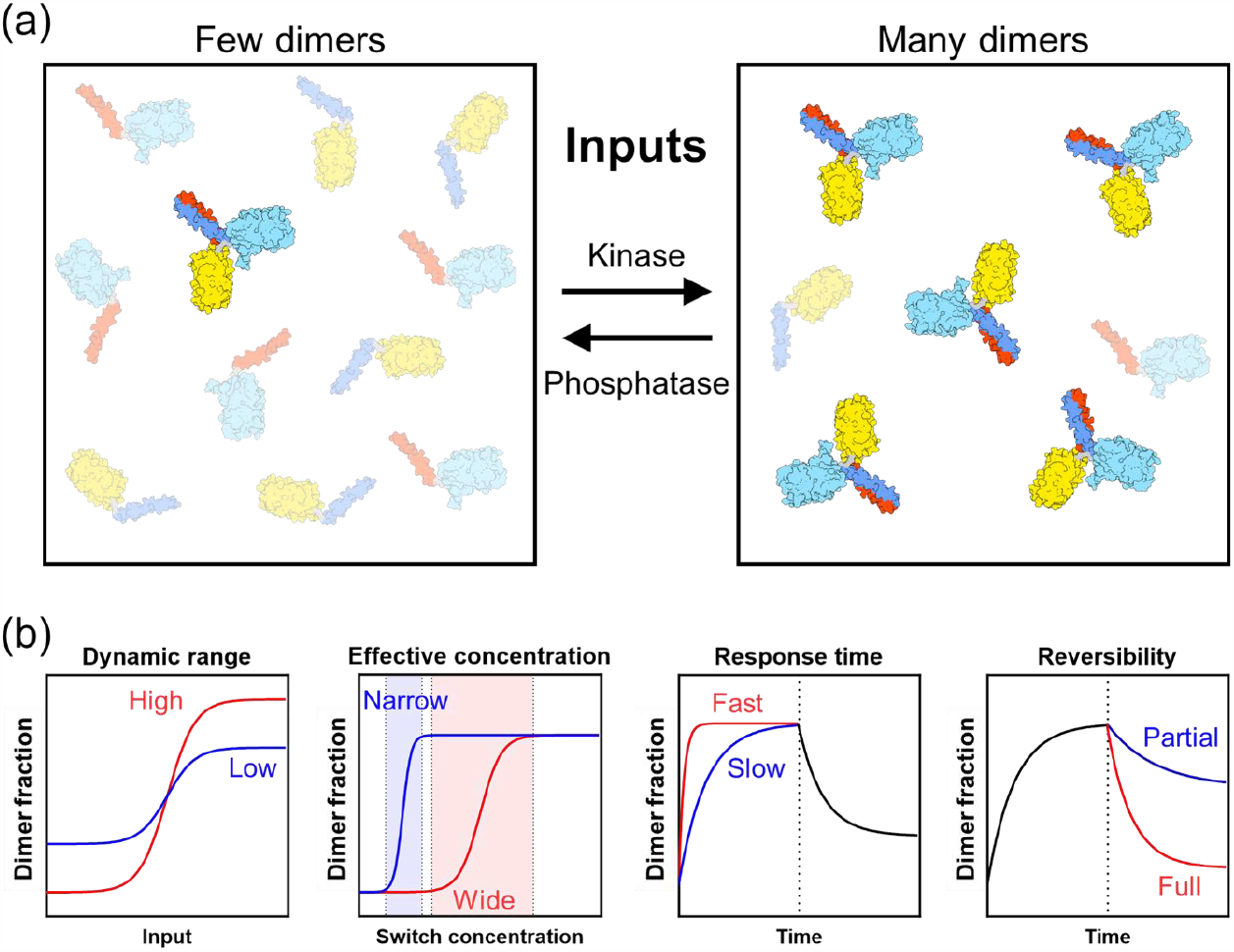
Performance metrics of a phosphorylation-driven protein switch. **(a)** Phosphorylation of a two-component protein switch by a protein kinase strengthens the inter-subunit interaction, increasing the concentration of active protein switch dimers. Conversely, dephosphorylation by a protein phosphatase shifts the equilibrium towards inactive switch monomers. Monomers are shown as transparent graphics to more easily visualize the number of depicted dimers **(b)** Illustrations of performance metrics for protein switches examined in the present study. The dynamic range and effective concentration metrics are steady-state equilibria of monomer vs dimer in respect to the input signal or protein switch concentration, whereas the response time and reversibility are kinetic metrics in respect to time. For the response time and reversibility graph, the vertical dotted line represents the changes in dimerization from phosphorylation of the protein switch (left of the dotted line) and from dephosphorylation of the protein switch (right of the dotted line).

Here, we introduce a theoretical framework to evaluate two-component protein switches, focusing on phosphorylation-responsive coiled coil interactions. Our models enable the evaluation of the four switch performance metrics: dynamic range, effective concentration, response time, and reversibility. Our framework combines the binding and phosphorylation kinetics of switches to predict their behavior across varying concentrations and time intervals. Using these insights, we design and evaluate two novel phosphorylation-driven protein switches and demonstrated their responsiveness to specific kinases and phosphatases. These switches feature phosphorylation-sensitive coiled coils fused to fluorescent proteins, enabling real-time measurements through Förster resonance energy transfer (FRET). Our model-guided design showcases the switches’ fast response time, reversibility, and wide dynamic range, confirming their potential as biosensors for practical applications.

## Results

### Binding equilibria delimit the dynamic range and effective concentration of phosphorylation-driven switches

Protein switches regulated by phosphorylation such as those that occur in nature or as engineered proteins involve a strengthening or weakening of the interaction between two PPI interfaces upon phosphorylation or dephosphorylation of specific residues at or near the binding interfaces. The interfaces of the PPI thus serve as sensor domains that can be fused to actuators to produce an output, for example, the halves of a split GFP (to produce a fluorescent signal output)^5^ or lipid-binding domains for cell membrane targeting^4^. At the molecular level, the addition of negatively charged phosphate groups to serine, threonine, or tyrosine residues by the activity of a kinase can create attractive or repulsive interactions between the interfaces, due to the addition or removal of negative charges and steric bulks. Changes in the conformation of individual components may also affect the strength of the interaction. Ultimately, the effect of protein phosphorylation on a two-component protein switch can be summarized as a change in the dissociation constant (*K*_*d*_) of the PPI, where lower *K*_*d*_ values correspond to stronger interactions. Given that *K*_*d*_ is one of the most fundamental descriptors of a given PPI, we modeled the effect that phosphorylation has on a protein switch as a function of the *K*_*d*_ of a PPI before and after phosphorylation.

The equation for the dissociation constant between two proteins that can form a heterodimeric complex (Equation 1) can be expanded to yield an expression that describes the concentration of the dimeric species as a function of the total concentration of each monomer and the *K*_*d*_ of the interaction (Equation 2). Using this expression, we can define several parameters that describe the steady-state properties of a phosphorylation responsive switch. Firstly, as the maximum dimer concentration is limited by the least concentrated monomer, the dimer fraction (*D*) is given in Equation 3. Further, the effect of phosphorylation on a PPI is defined as either the fold-change of the dimer fraction (*r*, Equation 4) or the absolute change (Δ, Equation 5) as a function of the total concentration of the monomers and the *K*_*d*_ values before and after addition of phosphate groups (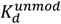 and 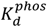) to the protein interfaces. Together, these equations can be used to describe the response of a protein switch to phosphorylation in terms of effective concentration range and dynamic range as a function of four key parameters: the total concentrations of the protein monomers ([*A*]_0_ and [*B*]_0_) and the *K*_*d*_ values of the PPI before and after phosphorylation (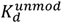 and 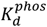).

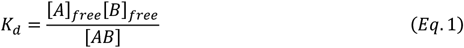

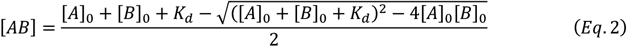

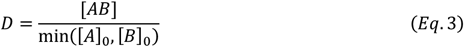

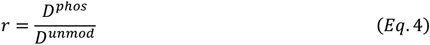

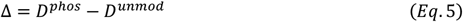

The above equations were used in a series of computational models to predict how total monomer concentrations and *K*_*d*_ values affect the behavior of phosphorylation-driven protein switches. The word “switch” implies the existence of an “off” state and an “on” state that can switch in response to stimuli (here, phosphorylation and dephosphorylation by kinases and phosphatases, respectively). At the single molecule level, this is intuitively true: two protein molecules are either interacting or they are not. However, functional applications of protein switches, such as within a cell, would be at concentrations where the protein molecules exist in an equilibrium of monomers and dimers (Figure 1a). An ideal protein switch would start exclusively as monomeric subunits and completely transition to dimers (or vice versa) in response to phosphorylation in a “all-or-nothing” fashion, which would require the *K*_*d*_ of the two protein components to be infinitely high before phosphorylation and equal to zero after phosphorylation, a physically impossible scenario. We therefore considered three dynamic ranges that approximate the ideal protein switch: a transition from 1% to 99% dimers (*r* = 99); from 10% to 90% dimers (*r* = 9); and from 25% to 75% dimers (*r* = 3). The model was used to calculate the *K*_*d*_ values required to realize these changes in dimerization in response to phosphorylation at a concentration of 1 μM of each subunit of the protein switch (the effect of changing the switch concentration is discussed further below). The predicted *K*_*d*_ values for each dynamic range are summarized in Figure 2 and Table 1. Our model predicts that to realize a 99-fold dynamic range in response to phosphorylation from 1% to 99% dimers at 1 μM concentration, the *K*_*d*_ value between the two components of the protein must switch from 9.8 × 10^-5^ M to 9.9 × 10^-11^ M, that is, from the high micromolar regime to the low picomolar regime. This corresponds to an extreme change in *K*_*d*_ of nearly one million-fold (or a change in free energy of dimerization of approximately 35 kJ/mol upon phosphorylation). To date, such extreme responses to phosphorylation have not been measured in natural proteins. For example, some of the strongest phosphorylation-dependent interactions exhibit *K*_*d*_ values in the nanomolar regime^14–18^ whereas in other cases, phosphorylation modulates the *K*_*d*_ of a protein interaction within the micromolar regime^19–21^.

**Table 1.** Predicted dissociation constants for a two-component protein switch to realize pre-defined dynamic ranges at a 1 μM concentration.

| Dimer fraction | | | Predicted $K_d$ values | | |
| --- | --- | --- | --- | --- | --- |
| Before phosphorylation | After phosphorylation | Fold-change | $K_d^{unmod}$ | $K_d^{phos}$ | Fold-change |
| 0.01 | 0.99 | 99 | $9.8 \times 10^{-5}$ M | $9.9 \times 10^{-11}$ M | 1/989,899 |
| 0.10 | 0.90 | 9 | $8.1 \times 10^{-6}$ M | $1.1 \times 10^{-8}$ M | 1/736 |
| 0.25 | 0.75 | 3 | $2.25 \times 10^{-6}$ M | $8.3 \times 10^{-8}$ M | 1/27 |

**Figure 2.**
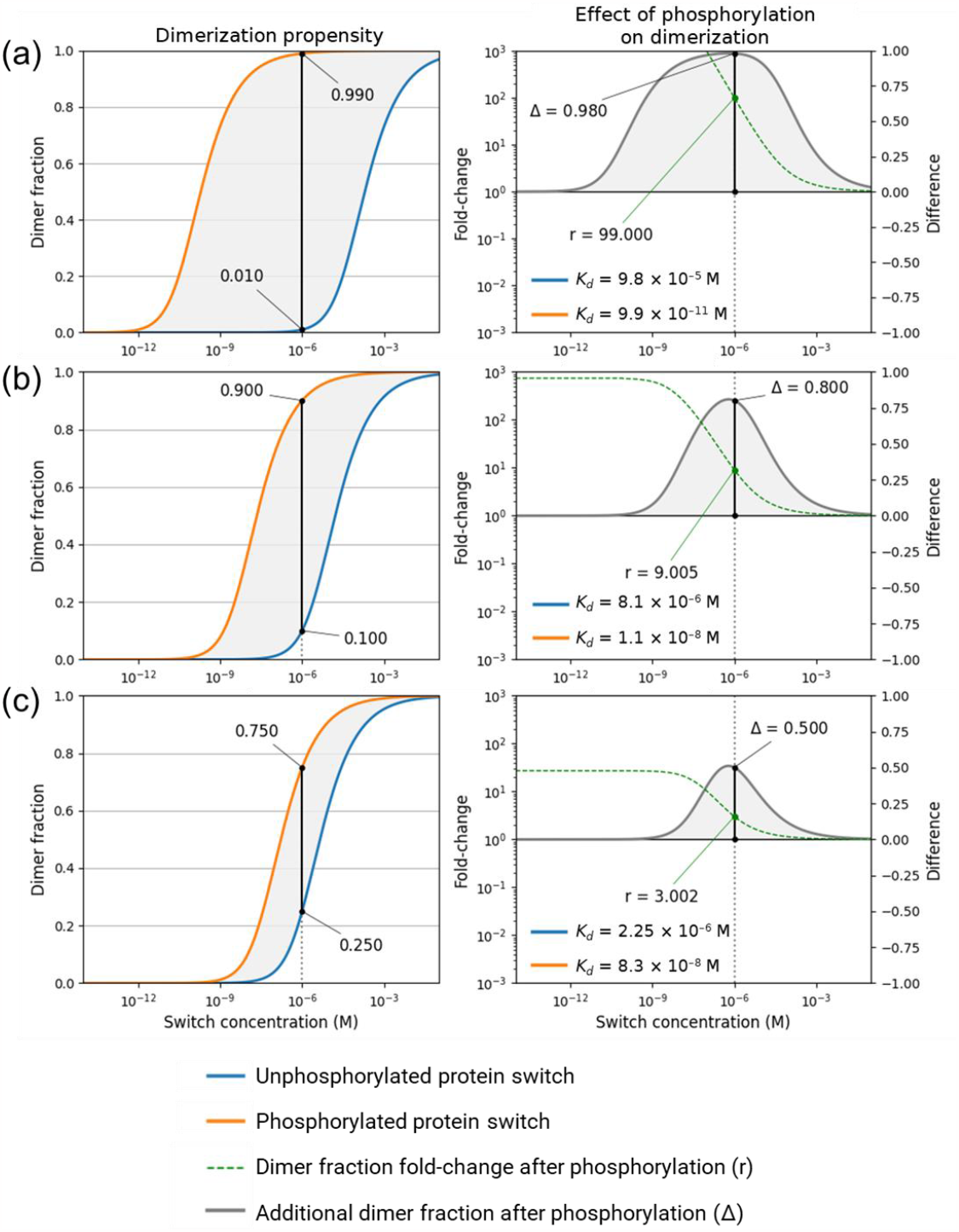
Predicted dimer fractions of a two-component protein switch in response to phosphorylation. Predicted K_d_ values of a protein switch at a 1 μM concentration that are required to achieve **(a)** a 99-fold dynamic range; **(b)** a 9-fold dynamic range; and **(c)** a 3-fold dynamic range. On the left, the dimer fraction as a function of the protein switch concentration for each dynamic range is plotted in blue for the unphosphorylated state and in orange for the phosphorylated state. A vertical black bar highlights the dynamic range at 1 μM protein switch. On the right, the relative change (see Equation 4) is plotted as a dashed green line and the absolute change in dimer fraction (see Equation 5) is plotted as a black line as a function of the protein switch concentration.

We also considered the effect of switch concentration on the steady state properties of our model systems to investigate their effective concentration ranges (the concentrations that produce a substantial fold-change in the dimer fraction upon phosphorylation). The modelling results suggest PPI-mediated protein switches display minimal dynamic range at picomolar and high millimolar concentrations as the binding equilibria tend to the complete dissociation or association, respectively, of the switch components. At these concentrations, PPI-mediated protein switches cease to be “switchable”. Regardless, greater differences between *K*_*d*_ values before and after phosphorylation produce a wider concentration range in which a protein switch retains its capacity to respond to phosphorylation. To further quantify this point, the dimer fractions before and after phosphorylation for each pair of *K*_*d*_ values listed in Table 1 were also calculated using protein switch concentrations of 10 nM and 100 μM (Table 2 and Figure S1). The predictions show that, in all cases, decreasing the concentration of the protein switch enhances the “signal-to-noise” ratio. That is, lowering the switch concentration decreases the dimer fraction before phosphorylation and increases the fold-change of the dimer fraction in response to phosphorylation. Increasing the protein switch concentration has the converse effect. This analysis reveals that, in fact, the dynamic range of a two-component protein switch tends to a value equal to 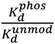 when the protein switch concentration tends to zero.

**Table 2.** Predicted dynamic ranges at different concentrations of protein switches.

| $K_d$ values | | | Predicted changes in dimer fractions <sup>a</sup> | | |
| --- | --- | --- | --- | --- | --- |
| $K_d^{unphos}$ | $K_d^{phos}$ | Fold-change | 10 nM switch | 1 μM switch | 100 μM switch |
| $9.8 \times 10^{-5}$ M | $9.9 \times 10^{-11}$ M | 1/989,899 | 0.000 → 0.905<br>($>10,000\times$) | 0.010 → 0.990<br>(99×) | 0.385 → 0.999<br>(2.6×) |
| $8.1 \times 10^{-6}$ M | $1.1 \times 10^{-8}$ M | 1/736 | 0.001 → 0.366<br>(366×) | 0.100 → 0.900<br>(9×) | 0.753 → 0.990<br>(1.3×) |
| $2.25 \times 10^{-6}$ M | $8.3 \times 10^{-8}$ M | 1/27 | 0.004 → 0.098<br>(24.5×) | 0.250 → 0.750<br>(3×) | 0.861 → 0.972<br>(1.1×) |

<sup>a</sup> Dimer fraction of the protein switch before and after phosphorylation, separated by an arrow, for each switch concentration. The numbers in brackets indicate the fold-change in dimer fraction.

However, as the fold-change (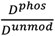, equation 4) in the dimer fraction increases with lower concentrations of the protein switch, the absolute change (*D*^*phos*^ − *D*^*unmod*^, equation 5) in dimer fraction becomes negligible and likely undetectable. Altogether, these results show an inherent trade-off between the baseline dimer concentration and the absolute response to phosphorylation (Figure 2). Thus, the intended application of a protein switch should be considered to determine the relevant protein switch concentrations, an acceptable baseline dimer concentration (noise) and the necessary dimer fold-change (*r*) or absolute change (Δ) upon phosphorylation to produce a measurable output (signal). Taken together, these considerations can be used to guide the choice of 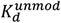 and 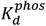 values when designing a protein switch.

In summary, the binding equilibria model highlights two parameters that can be modulated to tune the performance of a protein switch: the *K*_*d*_ values before and after phosphorylation of the protein switch and the concentration at which the protein switch is used. Together, these parameters control two important performance metrics of a protein switch: the dynamic range of the switch and the effective concentration of the protein switch, which is the concentration range at which the switch exhibits a response to phosphorylation.

### The interplay between phosphorylation and binding kinetics determines the reversibility of phosphorylation-driven switches

The performance of a protein switch can also be assessed via two time-dependent metrics: its response time to phosphorylation by a kinase and its reversibility after dephosphorylation by a phosphatase. The response time is the time taken for a protein switch to fully convert from the baseline dimer concentration to the maximum achievable dimer concentration. The reversibility of a protein switch refers to the capacity of the switch to revert from the dimeric state to the monomeric state after dephosphorylation, and how rapidly. Assessing the response time and reversibility of a switch requires consideration of PPI and PTM kinetics. The binding kinetics are determined by the association rate (*k*_*on*_) and dissociation rate (*k*_*off*_) between the two interacting subunits of the switch, with the two rates determining the dissociation constant (*K*_*d*_) used in the equilibria model (Equation 6). The PTM kinetics correspond to the rates of phosphorylation (*k*_*p*_) and dephosphorylation (*k*_−*p*_) of the protein subunits. For a two-component protein switch controlled by one phosphorylation site, the binding kinetics between its protein components (monomers A and B) will depend on the unphosphorylated and phosphorylated states of the switch, which requires four binding kinetics parameters 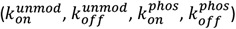. Furthermore, the activity of the kinase and phosphatase towards the monomeric or dimeric states of the switch may differ, for example if the phosphorylation site is occluded in the dimeric state. Thus, four additional phosphorylation kinetics parameters are required 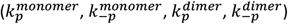. Together, the eight kinetic parameters (Figure 3a) can be incorporated into equations 7.1 to 7.5 to describe how the concentrations of the species *A, B, Ap, AB*, and *ApB* evolve in respect to time, where *p* denotes a single phosphorylation site on monomer *A* (Figure 3a). Note that for a protein switch modulated by two phosphorylation sites, such as peptide pairs described previously^3^, eight binding kinetics parameters and sixteen PTM kinetics parameters would be necessary to describe the entire system (Figure S2).

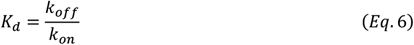

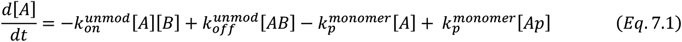

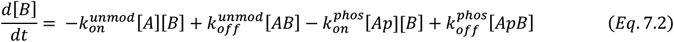

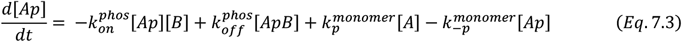

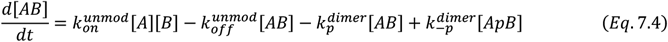

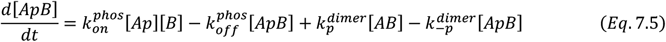

We numerically solved the above ordinary differential equations to investigate the dynamics of our model protein switches in response to phosphorylation. Specifically, our kinetic model comprises three stages. First, the initial concentrations of monomers and binding kinetic parameters are set, and the system is equilibrated to reach the baseline dimer concentration (in this stage, all PTM kinetics parameters are set to zero). Secondly, the phosphorylation kinetics parameters (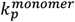 and 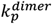) are updated to reflect the addition of kinase activity, and the system relaxes towards a new equilibrium as the protein switch is phosphorylated. Thirdly, the dephosphorylation parameters (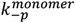 and 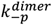) are updated to reflect the addition of phosphatase activity to the system that competes with the kinase. An example set of inputs for the kinetic model is shown in Figure 3b and the output is shown in Figures 3c and 3d. In this example, the *k*_*on*_ and *k*_*off*_ parameters are set to values that result in 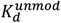 and 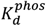 values of 2.25 × 10^-6^ M and 8.3 × 10^-8^ M (that were predicted for a 3-fold dynamic range; see Table 1), and the 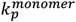 and 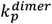 parameters were set to an arbitrarily chosen value of 10^-2^ s^-1^ to reflect the addition of a kinase that is equally active towards the monomeric and dimeric states of the switch. The kinetic model predicted a change in dimer fractions from 25% to 75% following phosphorylation for a protein switch concentration at 1 μM, in concordance with the equilibria model, but it also enabled to calculate the time required for the switch to attain the maximum response to the kinase input.

**Figure 3.**
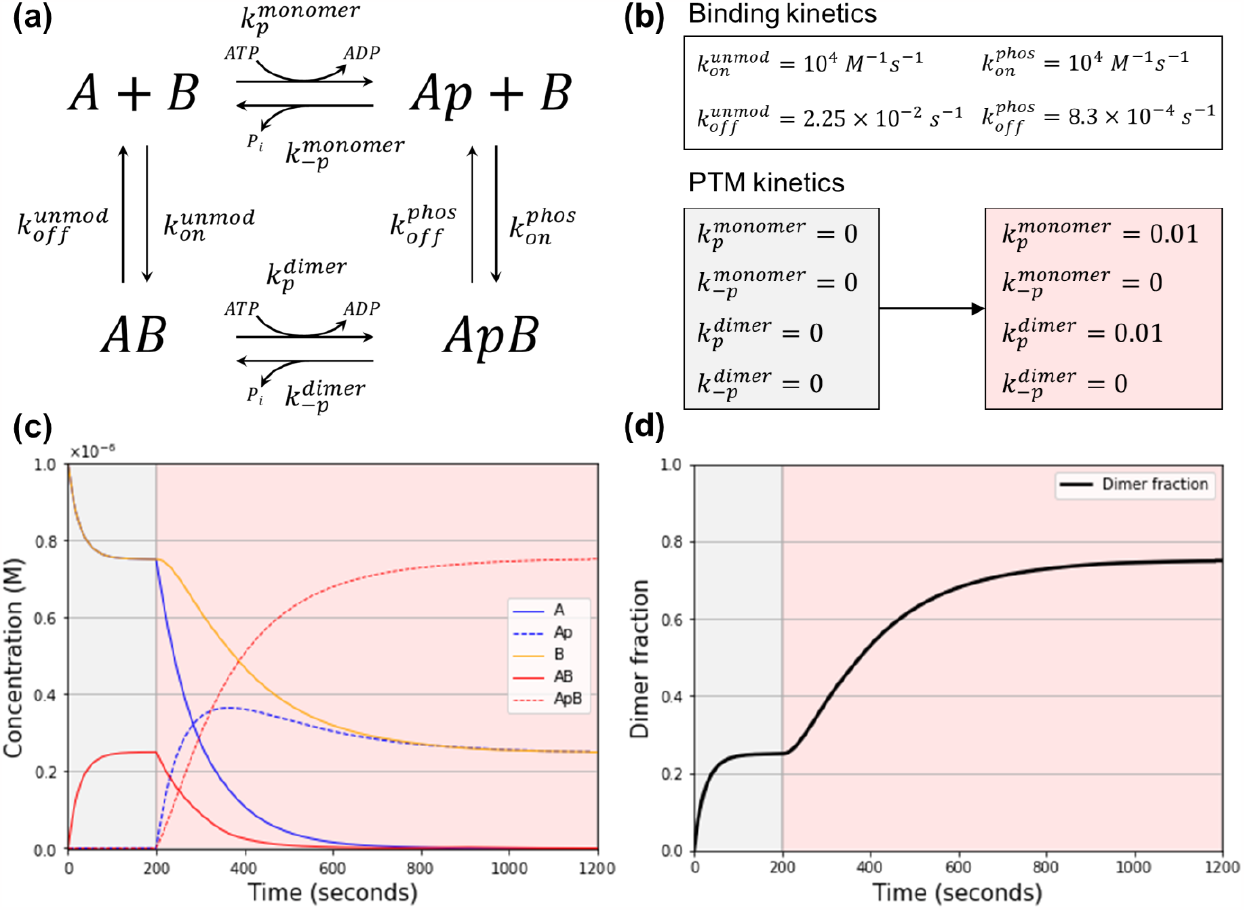
A kinetic model of a phosphorylation-driven protein switch. **(a)** A protein switch comprising two interacting monomers, whose interaction is modulated by a single phosphorylation site, can be described as the relative concentrations of five species: monomers A, B, and Ap, and dimers AB and ApB, where p denotes the single phosphorylation site on A. **(b)** Four binding kinetics parameters and four PTM kinetics parameters describe the evolution of each concentration in respect to time. The four kinetic parameters are updated between the equilibration (grey box) and the phosphorylation (red box) stages. **(c)** Evolution of the concentration of the five protein entities in the system during the equilibration (grey background) and phosphorylation (red background) stages of the simulation using the parameters from (b). **(d)** Same as in (c) but showing the evolution of the dimeric fraction of the system.

The model was used to evaluate how the binding kinetic parameters *k*_*on*_ and *k*_*off*_ influence the response time of a switch to the same amount of kinase activity. For any pair of *k*_*on*_ and *k*_*off*_ values that result in the same *K*_*d*_ value (see equation 5), the dimer concentration after phosphorylation is the same. However, as shown in Figure 4a, the exact *k*_*on*_ and *k*_*off*_ values influence the response time of the switch. In these simulations, 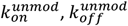, and 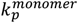 and 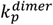 were constant, but the 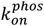 and 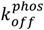 values (the binding kinetics after phosphorylation) were varied. For the same 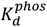 value of 8.3 × 10^-8^ M, the higher the values for 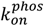 and 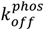, the faster the response rate, until a limit that is set by the overall phosphorylation rate (here, defined by 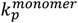 and 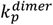). At this limit, the formation of new dimers evolves at the exact same rate as the addition of phosphate groups to the protein switch.

**Figure 4.**
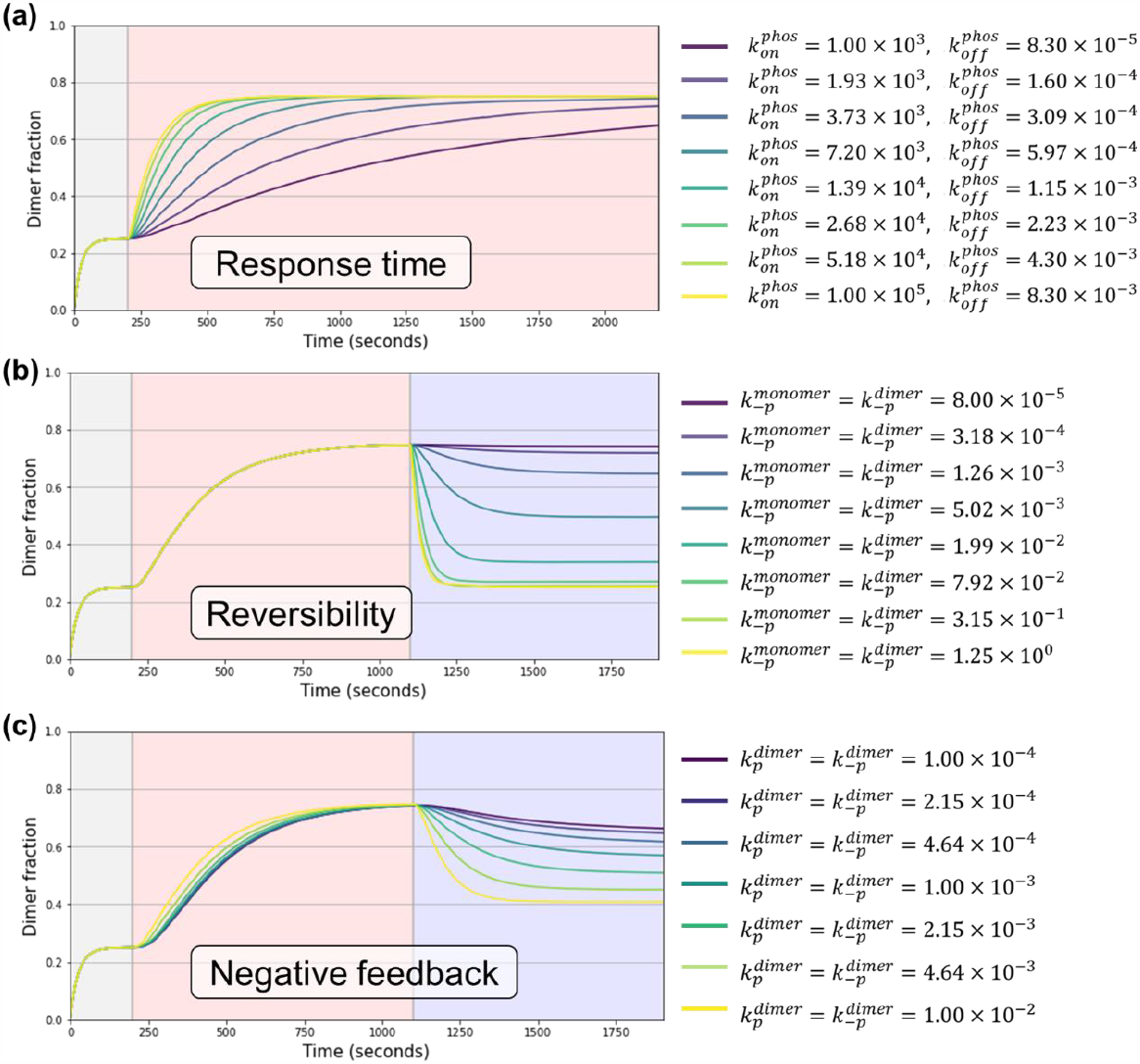
The interplay between binding and phosphorylation kinetics delimits protein switch performance. **(a)** For a same 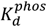 value following phosphorylation of the switch, increasing interaction turnover rates (increasing 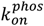 and 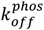 values, from purple to yellow graphs) are predicted to improve the response time of the switch. **(b)** Phosphatase activity must be significantly higher than kinase activity to revert the protein switch to its baseline dimer fraction. The graphs show the predicted reversibility of the switch for phosphatase activities ranging from 125 times lower (purple graph) to 125 higher (yellow graph) than the competing kinase activity. **(c)** Dimerization of a two-component protein switch in response to phosphorylation can render the switch less sensitive to inputs, affecting reversibility. The graphs show the predicted response of a switch whose dimerization does not affect (yellow graph) or greatly attenuates its likelihood to be (de)phosphorylated 100-fold (purple graph). The graph background colors indicate the equilibration (grey), phosphorylation (red), and dephosphorylation (blue) stages of the simulation; during the third stage of the simulation, the kinase activity persists and competes with phosphatase activity. The unit of k_p_, k_−p_, and k_off_ is s^-1^ and the unit of k_on_ is M^-1^s^-1^.

The reversibility of the protein switch was investigated using the kinetic model with simulations that included phosphatase activity (defined by the parameters 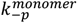 and 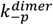) after the system had equilibrated in the presence of kinase activity. Kinase activity (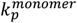 and 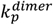) was retained in this simulation to represent the switch responding to both kinase and phosphatase activities simultaneously. The simulations show that a phosphorylation-driven switch can be reversed to the baseline dimer fraction when the phosphatase activity sufficiently exceeds the kinase activity, whereas phosphatase activities lesser than the kinase activity only slightly attenuate the phosphorylation-driven dimerization of the switch (Figure 4b). Starting with the same *k*_*on*_, *k*_*off*_, 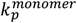, and 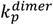 parameters as those shown in Figure 3b, we tested several dephosphorylation rates 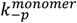 and 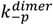 ranging from 125-fold higher to 125-fold lower than the phosphorylation rates 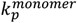 and 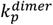. At *k*_−*p*_ = *k*_*p*_, the switch partially reverses to a dimer fraction of 0.41 (compared to the 0.25 baseline, before the addition of the kinase input signal). At *k*_−*p*_ = 2 × *k*_*p*_, the switch reverses to a dimer fraction of 0.34, and at *k*_−*p*_ > 5 × *k*_*p*_, the switch nearly completely reverses to dimer fractions of less than 0.29. The relationship between PTM kinetics and switch reversibility is plotted in Figure S3. The model assumes an infinite supply of ATP and does not account for ATP depletion over time. For *in vivo* applications, it can be assumed that ATP is continuously produced by the cell through respiration or fermentation. In contrast, in an *in vitro* system, the kinase would eventually run out of ATP, at which point the phosphatase would completely dephosphorylate the switch. Taken together, these simulations also challenge the concept of a “switch” that can be switched off as easily as it can be switched on.

The previous simulations assumed the kinase and phosphatase showed equal activities towards the monomeric and dimeric states of the protein switch (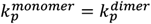 and 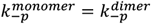). The next simulations examined whether a feedback loop could be formed between the activation of the switch and its sensitivity to the input signals. Specifically, we were interested in whether a negative feedback loop would result from the occlusion of the phosphorylation site upon dimerization of the switch. Structural data of phosphorylation-mediated PPIs has shown previously that the phosphorylated residue on a protein interface can be occluded when bound by its binding partner,^22,23^ and thereby prevent the access of protein phosphatases and inhibit dephosphorylation^14^. We thus considered systems where the dimeric state of the switch could still be phosphorylated or dephosphorylated, but at slower rates than the monomeric states (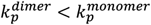 and 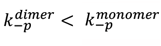). The simulations showed that competition between dimerization and phosphorylation events has a minor impact on the response to the kinase input, with slightly slower response to phosphorylation if there is a negative feedback loop (Figure 4c). However, the effect on the reversibility of the switch was pronounced. In this system, the switch primarily exists in the monomeric state at first, which is more sensitive to phosphorylation signals. Addition of the phosphatase results in primarily a dimeric state, which is less sensitive to dephosphorylation signals, and the response time to the phosphatase is lengthened. We also investigated whether varying the protein interaction turnover rate (varying the *k*_*on*_ and *k*_*off*_ values while maintaining the *K*_*d*_ values constant) could affect reversibility. The simulations predict that faster interaction turnover rates in the phosphorylated state (higher 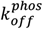 and 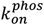) exacerbate the “negative feedback” effect, rendering the protein switch nearly irreversible (Figure S4). Conversely, slower interaction turnover rates in the phosphorylated state (lower 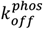 and 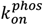) vastly improved the sensitivity of the switch to phosphatase activity, which improved its reversibility and compensated for the negative feedback loop. Moreover, the simulations predicted an intriguing behavior of a protein switch with slow interaction turnover rates: upon the addition of kinase activity, the system is perturbed with a temporary decrease in dimer concentration before the dimer fraction increases in response to phosphorylation, as is expected from the switch (Figure S4c), akin to one oscillation. This behavior is counter-intuitive, but it is relevant for applications where oscillations of a protein switch cannot be tolerated. Finally, these additional simulations reveal an important trade-off: as shown in Figure 4a, faster interaction turnover rates in the phosphorylated state improve the response time of the protein switch. Thus, there exists an inherent trade-off between response time and reversibility of a protein switch.

Altogether, the kinetic model of phosphorylation-driven switches reveals an intricate relationship between binding and PTM kinetics. The response time of a protein switch is intimately linked to the interaction turnover rate (the *k*_*on*_ and *k*_*off*_ values that compose the *K*_*d*_) in two ways: first, slow turnover rates result in a delayed response to phosphorylation because the switch transitions from the monomeric to the dimeric state more slowly than the rate at which it is phosphorylated. Second, a slow turnover rate may delay phosphorylation itself if one state (monomeric or dimeric) is the better substrate for the kinase or phosphatase. In addition, a switch is never truly fully reversible, unless the system allows the complete removal of the kinase input signal before the addition of a phosphatase.

### Design of phosphorylation-driven protein switches using coiled coils as sensor domains

Protein switches were designed and recombinantly produced for use in validating the kinetic model, for example, the effective concentration range for a switch to transition from monomer to dimer upon an input signal. The switches were based upon three coiled coil-forming peptides from our previously published work^3^, named E3-Switch, E3_A_-Switch, and K3-Switch. The E3-Switch and K3-Switch are based on the well characterized E3/K3 heterodimeric coiled coil^24^ with the introduction of one PKA recognition site in each helices of the coiled coil, judiciously placed to promote attractive charged interactions upon phosphorylation of the target serine, thereby stabilizing the interaction. E3_A_-Switch is similar to E3-Switch, except a key leucine residue is mutated to alanine, to modulate the interaction strength between the two peptides. Combining either E3-Switch or E3_A_-Switch with K3-Switch was shown to form dimeric coiled coils, with the E3/K3-Switch and E3_A_/K3-Switch having *K*_*d*_ values of 1.39 × 10^-8^ M and 3.2 × 10^-7^ M, respectively^3^. Importantly, the interactions between the peptides were strengthened upon phosphorylation by PKA with an observed decrease in *K*_*d*_ values for the E3/K3-Switch (an undetermined subnanomolar value) and the E3_A_/K3-Switch (6.7 × 10^-8^ M). Furthermore, λPP could be used to dephosphorylate the peptides and reverse the increased affinity.

The coiled coil-forming peptides were genetically fused to the C-terminus of either mCerulean3 (CFP) or mVenus (YFP) via a TEV protease cleavage sequence to create the three proteins CFP-E3-Switch, CFP-E3_A_-Switch, and YFP-K3-Switch (Figure 5a and Table S1). CFP and YFP can engage in Förster resonance energy transfer (FRET) when < 5 nm in proximity, whereby excitation of CFP at 433 nm results in energy transfer to YFP and emission at 533 nm (Figure 5b). The FRET pair was chosen as the actuating domains of the switches because FRET is a nanosecond phenomenon, and thus well suited for real time measurements of protein interaction dynamics. In addition, the FRET output can be used to measure the *K*_*d*_ of the interacting peptides in response to changes in phosphorylation. We hypothesized that the CFP-E3-Switch/YFP-K3-Switch and CFP-E3_A_-Switch/YFP-K3-Switch pairs (hereafter, simply E3/K3-Switch and E3_A_/K3-Switch) will function as phosphorylation-driven protein switches with FRET as the output signal, because the strength of the interaction between the coiled coils would be modulated by PKA-mediated phosphorylation and λPP-mediated dephosphorylation.

**Figure 5.**
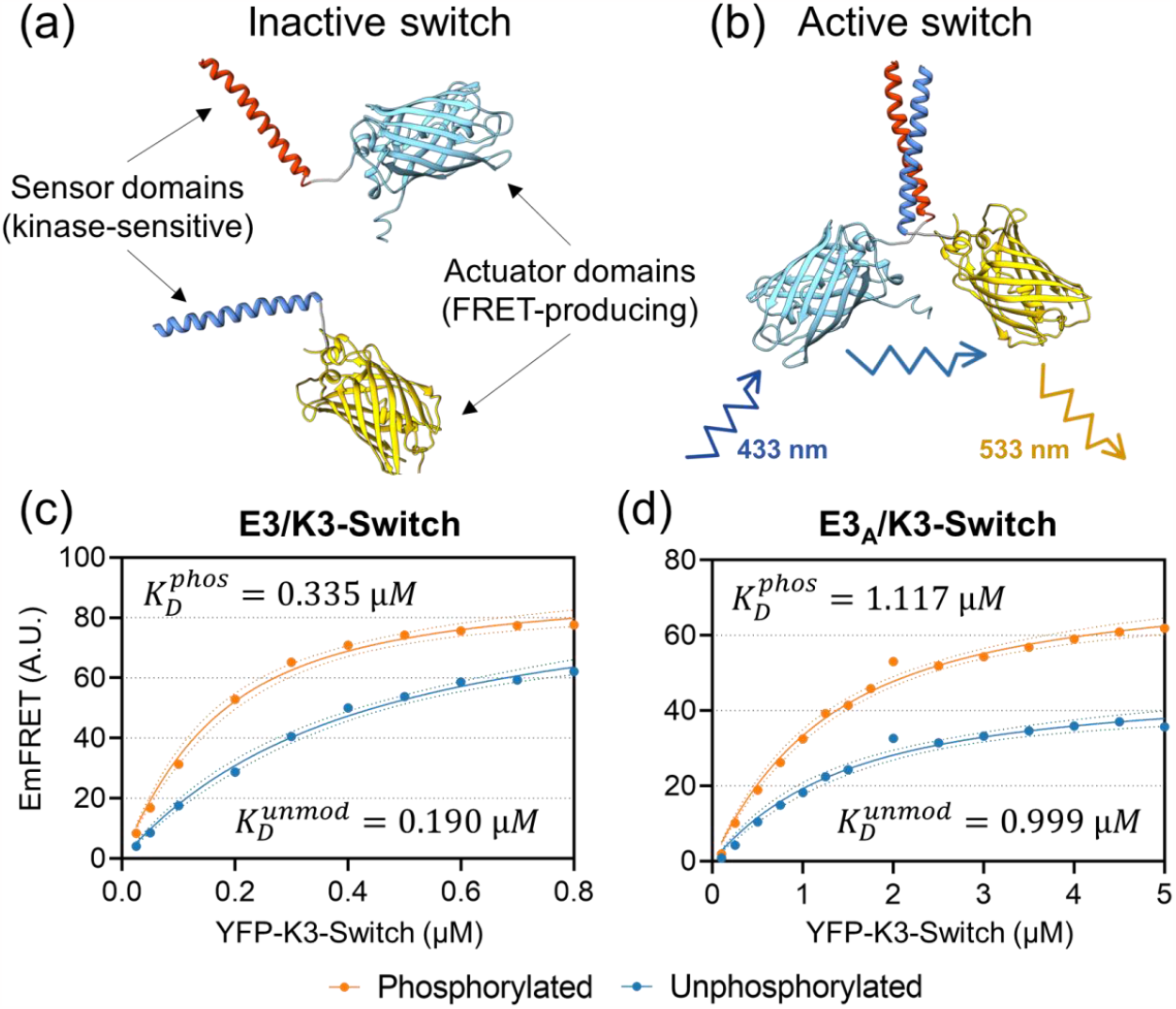
Dissociation constant and response of E3/K3-Switch and E3_A_/K3-Switch calculated from FRET measurements. **(a, b)** Design of a dimeric protein switch where each component is a fusion between a phosphorylatable peptide (blue and red alpha-helices) to a fluorescent protein of a FRET pair (mCerulean3 and mVenus, in cyan and yellow, respectively). In the inactive state (a), the two components do not interact via the sensor domains and FRET cannot be produced upon excitation of the FRET donor, mCerulean3. In the active state (b), the two sensors interact as a heterodimeric coiled coil, bringing the actuator domains into proximity, enabling FRET. **(c)** Dissociation constants of E3/K3-Switch and **(d)** of E3_A_/K3-Switch in their non-phosphorylated (blue) and phosphorylated (orange) states measured with a FRET assay.

The recombinant proteins were expressed in *E. coli* cells and purified by immobilized metal affinity chromatography (IMAC). Native mass spectrometry confirmed that at a 10 μM concentration, the CFP-E3-Switch and YFP-K3-Switch were monomeric until combined, with the resulting E3/K3-Switch shown to be heterodimeric (Figure S5). The lack of self-interaction even at high 10 μM concentrations enabled application of the binding and kinetic models, which are based on heterodimeric binding events, to the study of our novel protein switches.

The *K*_*d*_ values of E3/K3-Switch and E3_A_/K3-Switch were measured before and after phosphorylation (Figure 5c and 5d, Table 3) using a FRET-based assay adapted from Song *et al*.^25^. CFP-E3-Switch or CFP-E3_A_-Switch were mixed with increasing concentrations of YFP-K3-Switch in the presence or absence of PKA and the FRET emission was recorded for each mixture. Nonlinear regression analysis of the FRET emission as a function of YFP-K3-Switch concentration determined the E3/K3-Switch having a *K*_*d*_ of 3.35 × 10^-7^ M and 1.09 × 10^-7^ M before and after phosphorylation, respectively (Equation 8, where *C* is the concentration of the FRET donor and *Y* is the concentration of the FRET acceptor). The predicted maximum FRET emission was the same for both phosphorylation states (92 A.U.), indicating that the increase in FRET signal upon phosphorylation by PKA was due to an increase in the formation of dimers between CFP-E3-Switch and YFP-K3-Switch. For the E3_A_/K3-Switch, the calculated *K*_*d*_ values were 1.17 × 10^-6^ M and 9.99 × 10^-7^ M before and after phosphorylation. The decrease in *K*_*d*_ upon phosphorylation is modest, however the data (Figure 5d) shows that E3_A_/K3-Switch emits substantially more FRET signal upon phosphorylation, with maximum FRET emissions for E3_A_/K3-Switch differing between the unphosphorylated and phosphorylated states (47 and 76 A.U., respectively). The increase in maximum FRET suggests that a mechanism other than the formation of dimers contributed to the phosphorylation-driven increase in FRET signal, possibly a structural change of the dimer that places CFP and YFP in closer proximity or in a relative orientation that favors FRET between the two proteins^26–28^. For comparison, Figure S6 shows a simulated dataset where the FRET response of E3_A_/K3-Switch to PKA activity is entirely driven by the *K*_*d*_ values of 1.17 × 10^-6^ M and 9.99 × 10^-7^ M but the maximum FRET emission is the same for both phosphorylation states (47 A.U.).

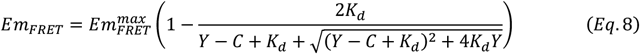

The equilibria model was used to estimate the effective concentration range for both switches. Using the *K*_*d*_ values from Table 3 as input for the model, the theoretical optimal concentrations in terms of fold-change and absolute change in the dimer fraction were calculated for E3/K3-Switch and E3_A_/K3-Switch (Figure S7). The model predicts an optimal concentration of 2.35 × 10^-7^ M for E3/K3-Switch and 1.31 × 10^-6^ M for E3_A_/K3-Switch to achieve the largest increase in dimer fraction following phosphorylation. However, as noted above, the relative change in dimer fraction can be improved by reducing the concentration of the protein switch (Table 2). To challenge the model, we prepared serial dilutions of equimolar mixtures of the proteins for the E3/K3-Switch and the E3_A_/K3-Switch and measured their FRET emission before and after addition of PKA (Figure S8). Following excitation at the CFP maximum of 433 nm, the ratiometric FRET (rFRET) can be reported as the ratio between the YFP maximum emission at 533 nm and the CFP maximum emission at 476 nm. As predicted by our model, the E3/K3-Switch exhibited an increase in rFRET response following phosphorylation as protein switch concentration decreases. In contrast, the optimal rFRET response for E3_A_/K3-Switch occurs at a concentration around 1 × 10^-6^ M, which is precisely the predicted concentration for the greatest absolute change in dimer fraction (Figure S7). No additional improvements of the fold-change of rFRET in response to phosphorylation were observed by lowering the concentration of the E3_A_/K3-Switch. The difference in protein switch behavior can be explained from the *K*_*d*_ measurements data (Figure 5c and 5d, Table 3): the response of E3/K3-Switch to PKA activity is driven primarily by changes in dimer concentrations, and therefore the equilibria model is well suited to predict its behavior. In contrast, E3_A_/K3-Switch exhibits only a minor decrease in *K*_*d*_ upon phosphorylation, but a substantial increase in FRET. The molecular mechanism underpinning this behavior is likely a phosphorylation-driven structural reconfiguration which, unlike PPI mechanisms, is insensitive to the total concentration of the protein switch. Thus, the equilibria model cannot fully predict the behavior of E3_A_/K3-Switch, and an updated model considering conformational equilibrium constants or conformational kinetics would be required to describe E3_A_/K3-Switch.

**Table 3.**
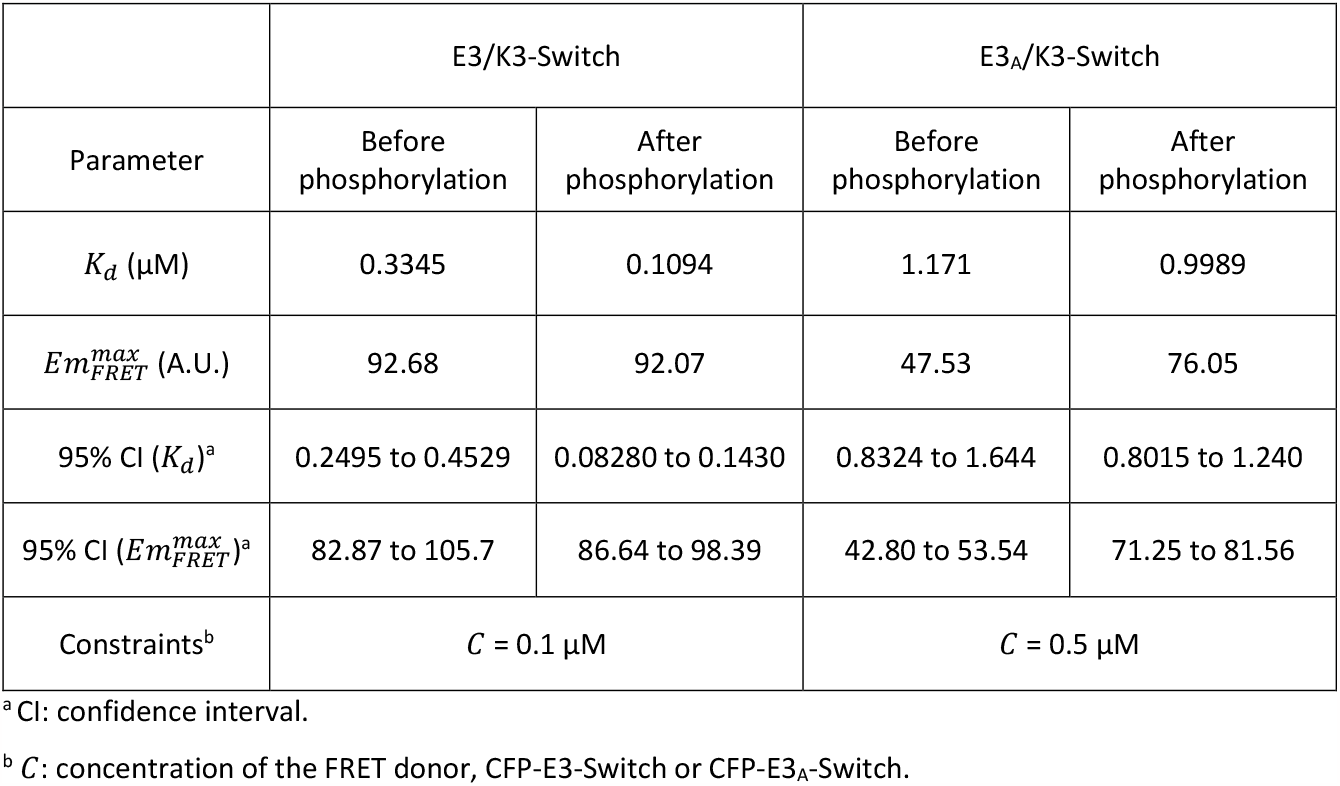
Calculated parameters from nonlinear regression analyses of FRET data.

Altogether, we successfully built novel phosphorylation-driven protein switches by appending a FRET protein pair as actuator to kinase-sensitive coiled coils as sensors. The qualitative performance of the switches could be inferred from the behavior of the coiled coils studied in isolation^3^, although the exact *K*_*d*_ values of the protein switch differed from those of the coiled coils alone, which is likely due to the fusion of the coiled coils to large protein subunits, and the differences in methodology employed for *K*_*d*_ measurement. Further, the detailed characterization of the protein switches using FRET as an output, combined with predictions from our models, revealed additional conformational mechanisms that underpin the response of the switches to PKA activity.

### Dynamic response of E3_A_/K3-Switch to kinase and phosphatase input signals

The real time response of E3/K3-Switch and E3_A_/K3-Switch to the activities of PKA and λPP were measured for comparison with the kinetic model. Prior to these experiments, we revisited the binding equilibria model to optimize the dynamic range of the two protein switches. In all previous experiments and simulations, we assumed that the concentration of the two components of the protein switch were equal. However, in FRET, the output signal is produced by the fraction of the FRET donor (here, CFP) that is in proximity to the FRET acceptor (YFP). Therefore, rather than considering the dimeric fraction in respect to the limiting monomer concentration (see Equation 3), one must consider the dimeric fraction in respect to the FRET donor concentration, and whether the donor or acceptor is the limiting monomer (Equation 9, where *A* is the donor, and *B* the acceptor).

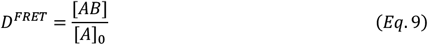

Using this definition of the FRET-producing dimer fraction, the binding equilibria model predicts that the dynamic range of phosphorylation-driven switches can be further improved when the acceptor concentration is higher than the donor concentration when the protein subunits concentration are in the vicinity of 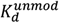 and 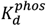 (Figure S9). Supporting these predictions, in the *K*_*d*_ measurement experiments, a more substantial increase in FRET was observed upon phosphorylation for data points where the concentration of the YFP acceptor was greater than the CFP donor (Figure 5c and 5d). Based on these observations and preliminary experiments (data not shown), we chose to work with 0.1 μM CFP-E3_A_-Switch 1 μM YFP-K3-Switch concentrations for the measurements of a protein switch performance in real time.

The E3_A_/K3-Switch was subjected to increasing concentrations of PKA (Figure 6a) and the rFRET was recorded in real time immediately following addition of recombinant PKA. As predicted, increasing the concentration of input kinase led to faster increases in rFRET. The initial velocity (*ν*_0_) was calculated by nonlinear regression using Equation 10, where *η* is the relaxation factor that accounts for nonlinearity in the system^29^. Although the reaction contains two PKA substrates (E3_A_-Switch and K3-Switch), the system can be simplified as the concentration of K3-Switch is tenfold greater than that of E3_A_-Switch, allowing us to approximate the increase in rFRET to a first order reaction. The initial velocity of the reactions correlated linearly with lower concentrations of PKA, indicating that the E3_A_/K3-Switch could be used to measure the activity of PKA through a simple FRET assay (Figure 6b). Overall, the reaction proceeded rapidly, with substantial FRET signal produced within 5 minutes, suggesting a fast protein interaction turnover between the switch subunits when comparing the FRET data to the simulations in Figure 4a.

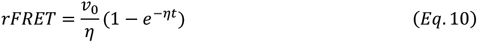

The reversibility of E3_A_/K3-Switch was measured by phosphorylating the switch using PKA then incubating the switch with varying amounts of λPP and measuring the rFRET over time (Figure 6c). Dephosphorylation buffer was added to the phosphorylated switch as a control, which caused a small decrease in FRET due to either the dilution of the samples or the interference of the buffer reagents with the protein switch dimerization. As expected, increasing the amount of λPP led to a more substantial decrease of the FRET signal. However, for the tested concentrations of λPP, the FRET signal did not revert to the baseline observed in (Figure 6a). When comparing the *in vitro* results to the simulations in Figure 4b, the FRET data suggest that the phosphatase activity was several-fold lower than the kinase activity present in the reaction. The E3_A_/K3-Switch is reversible but, as theory predicts, requires a vast excess of phosphatase activity to be fully deactivated in the presence of kinase activity.

**Figure 6.**
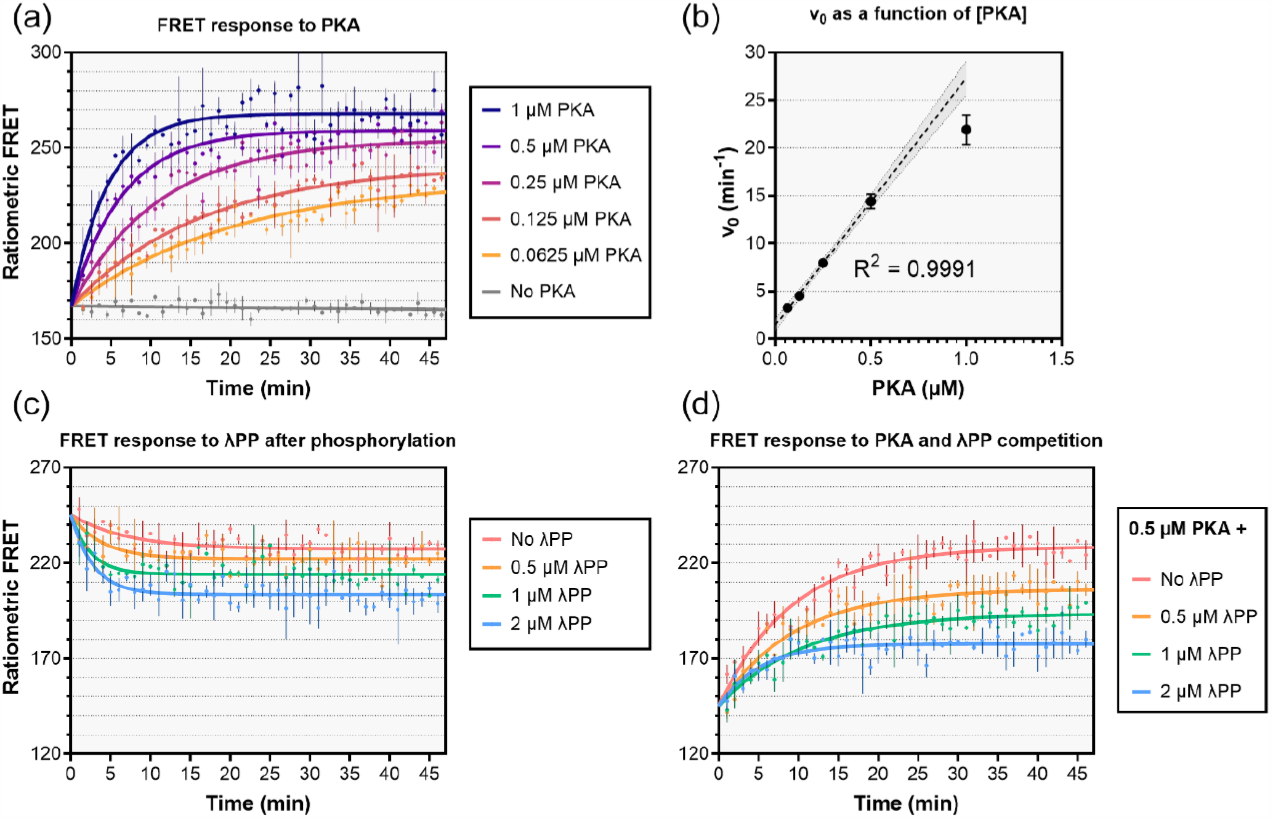
Real time response of E3_A_/K3-Switch to kinase and phosphatase activities. **(a)** Real time measurement of the rFRET emission of E3_A_/K3-Switch in response to several concentrations of PKA. **(b)** The initial velocities calculated by non-linear regression from the data in (a) plotted as a function of PKA concentration. Error bars correspond to the standard error of the best fit values of ν_0_. The linear regression for the first four data points is plotted as a dashed line. The shaded area corresponds to the 95% confidence interval. **(c)** Real time measurement of the rFRET emission of E3_A_/K3-Switch in response to several concentrations of λPP after phosphorylation of the protein switch. **(d)** Real time measurement of the rFRET emission of E3_A_/K3-Switch in response to a constant concentration of PKA against varying backgrounds of phosphatase activity. In (a), (c) and (d), data points represent the mean of two replicates and the error bars represent the standard deviation of the mean.

Finally, we subjected the E3_A_/K3-Switch to varying backgrounds of phosphatase activity prior to the addition of PKA (Figure 6d). This experiment, where kinase activity competes with phosphatase activity, reflects the conditions within living cells where phosphoproteins are constantly subject to and “integrate” phosphorylation and dephosphorylation signals ^30,31^. Real time rFRET revealed that a set PKA concentration produced different rates of FRET increase depending on the phosphatase background. Higher levels of phosphatase background activity resulted in slower rates of FRET increase, demonstrating that the E3_A_/K3-Switch could simultaneously respond to kinase and phosphatase inputs, sensing the balance of phosphorylation and dephosphorylation activities.

## Discussion

We developed a theoretical framework to evaluate PPI-driven protein switches in respect to four key performance metrics: dynamic range, effective concentration, response time, and reversibility. The first two metrics can be predicted based on binding equilibria equations, whereas predicting the two latter metrics requires several PTM and binding kinetic parameters. Our study focused on *de novo-* designed two-component switches that sense kinase and phosphatase activities as a model system. However, our modelling approach can be adapted to the study of natural proteins whose PPIs are modulated by PTMs or other types of biochemical or biophysical inputs that can affect the dissociation constant between the two subunits of a dimeric protein complex. Our equilibria and kinetic models enable the identification of potential applications of a protein switch with the calculation of its realizable dynamic ranges and response times. Moreover, the models revealed the existence of trade-offs between some performance metrics; for example, a trade-off between the absolute increase in the dimer fraction in response to kinase activity and the baseline dimer concentration (Table 2) and a trade-off between response time and reversibility (Figure 4 and Figure S4). In addition, we designed and produced two novel phosphorylation-driven protein switches that were characterized using our theoretical framework. Overall, our protein switches performed as predicted by the model and function as a tool to individually measure kinase or phosphatase activity in real time or integrate competing kinase and phosphatase activities.

Nonetheless, our theoretical and experimental work emphasizes important factors that are seldom considered in the design and function of two-component protein switches. First, the activation of dimeric switches such as those described in our and others’ previously published work is generally ascribed to a change in *K*_*d*_ between the two subunits upon phosphorylation. Here, the analyses of E3/K3-Switch and E3_A_/K3-Switch revealed additional factors that contribute to the performance of a switch. Although the FRET response of E3/K3-Switch to phosphorylation could be explained by the change in *K*_*d*_ between the subunits of the two protein switches (Figure 5a and 5b), our model underestimated the FRET response of E3_A_/K3-Switch. Specifically, our model is suitable for the description of changes in dimerization via the phosphorylation-sensing domains of the switch but does not account for phosphorylation-driven conformational changes or peculiarities of the actuating domains that may contribute to the response signal. In the case of FRET, the proximity and relative orientation of the FRET pair may be affected by the conformation of the sensor domains in addition to their dimerization. These results concur with surface plasmon resonance studies where the binding kinetics of certain dimeric coiled coils could only be modelled with the inclusion of a rate-limiting conformational rearrangement step^32^. Although our equilibria and kinetic models were created to be as generalizable as possible, variations of the models tailored to specific protein switch designs could include additional parameters, such as folding or conformational kinetics. Our data highlight the importance of characterizing the complete protein switch and not only the sensor domains in isolation, as has been done for *de novo* synthetic peptides that are not fused to actuating domains, as the properties of the sensor domains may not be fully transferred to a protein switch depending on the choice of actuators.

The choice of actuators may also impact the reversibility of the protein switch. In our new designs, we employed a pair of fluorescent proteins as FRET actuators that resonate at the electromagnetic level but do not otherwise physically interact and thus do not participate in the dimerization of the protein switch. In contrast, a previously described kinase-responsive protein switch employed split GFP as an actuator^5^. Split GFP comprises two subunits which, when brought into proximity, fold onto each other and irreversibly react to form the GFP chromophore. Thus, while the authors could demonstrate that phosphorylation of the sensor domains could be removed with a protein phosphatase, the fluorescence signal emitted by the actuating domains could not be attenuated after activation of the switch. Our choice of a FRET pair instead of split GFP to produce a fluorescent signal in response to phosphorylation resulted in a truly reversible switch, within the limits stipulated by our kinetic model (Figure 4b), where the output signal reflected the presence of phosphatase activity (Figure 6a and 6b).

Finally, our theoretical framework pays special consideration to an all-too-often neglected parameter: the concentration of the protein switch itself. This parameter underpins the dynamic range of the protein switch response to an input signal. For *in vitro* applications, such as the use of a protein switch as a biosensor to measure enzyme activity, it is relatively straightforward to tune the concentrations of each subunit and maximize dynamic range (Figures 5 and 6). Going forward, *in vivo* applications of phosphorylation-driven protein switches will likely attract more attention, including methods to deliver protein switches into living cells, either directly or encoded in nucleic acids. Controlling protein switch concentration and localization inside living cells will pose a challenge. For example, a tyrosine phosphorylation-responsive switch has been designed that operates inside mammalian cells to sense aberrant tyrosine protein kinase activity, indicative of a cancerous state, and to respond with the initiation of cell death^33^. However, the initial design resulted in an overly sensitive switch unable to distinguish between kinase activities *in vivo* and promoting the death of all cells, and several rounds of redesign were necessary to tune the switch. In the context of *in vivo* applications, which include diagnosis, therapeutics, and the design of synthetic signaling pathways, our theoretical framework should serve as a powerful tool to guide the design of phosphorylation-driven protein switches that operate at concentrations realizable within living cells.

## Materials and methods

### Binding equilibria and kinetic models of phosphorylation driven switches

Steady-state and kinetic models were implemented in Python using the SciPy library as per the equations described throughout the manuscript. The Matplotlib library was used to visualize the results. The scripts are available upon request.

### Construction of DNA plasmids and bacterial strains

DNA sequences encoding the E3-Switch-mCerulean3, E3_A_-Switch-mCerulean3, K3-Switch-mVenus, and λPP were codon-optimized for *E. coli* and synthesized by Integrated DNA Technologies with additional overhangs for Gibson assembly. The plasmid pET-19b (Novagen) was double digested using NcoI and BamHI and Gibson assembly performed to insert a gene construct into the plasmid. 50 μL of competent *E. coli* NEB Turbo cells (New England Biolabs) were transformed with each of the Gibson products by heat shock and selected on lysogeny broth (LB) agar plates (10 g/L sodium chloride, 10 g/L tryptone, 5 g/L yeast extract, 15 g/L agar) supplemented with ampicillin (100 μg/mL). Individual colonies from the transformation were inoculated into liquid LB growth medium (10 g/L sodium chloride, 10 g/L tryptone, 5 g/L yeast extract) complemented with ampicillin (100 μg/mL), grown overnight at 37 °C, and plasmid DNA extracted using the Monarch Plasmid Miniprep Kit (New England Biolabs), and Sanger sequencing used to confirm correct insertion of coding sequences at the Ramaciotti Centre for Genomics (UNSW Sydney). The pET-19b plasmid encoding PKA has been described in our previous work^3^.

### Protein expression and purification

Competent T7 Express *E. coli* cells (New England Biolabs) were transformed with either of the five plasmids encoding CFP-E3-Switch, CFP-E3_A_-Switch, YFP-K3-Switch, PKA, or λPP, and transformants were selected on LB agar plates complemented with ampicillin. Subsequently, a colony for each strain was used to inoculate 5 mL LB-ampicillin starter cultures, and the cultures were subsequently used to inoculate 200 mL of LB complemented with ampicillin in 1 L baffled flasks at a starting optical density of OD_600_ = 0.1. The cultures were incubated with shaking (200 RPM) at 37 °C until the OD_600_ was 0.6, and protein expression was induced with the addition of isopropyl-ß-D-1-thiogalactopyranoside (IPTG) to a final concentration of 1 mM. Protein expression was performed at 25 °C for 16 h, and the cells pelleted at 5,000 *g* for 10 min. Cell pellets were lysed by French pressing and the lysates clarified by centrifugation at 15,000 *g* followed by filtration using 0.22 μm Millex syringe filters (Merck). Protein purification was performed with an ÄKTA start chromatography system (GE Healthcare Life Sciences) using a 5 mL HisTrap FF column (GE Healthcare Life Sciences) as previously described^3^ except that an optimized buffer gradient was employed consisting of a first wash step of 5 column volumes (CV) at 0% elution buffer, a second wash step of 5 CV at 5% elution buffer, and a single elution step of 10 CV at 25% elution buffer. 5 mL fraction were collected and the fractions containing the recombinant protein at sufficient purity as assessed by SDS-PAGE were concentrated to approximately 5 mg/mL using Amicon Ultra-4 Centrifugal Filters with a 30 kDa molecular weight cut-off (MWCO) and stored at 4 °C with the addition of glycerol to a final concentration of 10% (v/v) and sodium azide to a final concentration of 0.05% (w/v). Depending on the requirements of subsequent experiments, proteins were either diluted directly into the final assay buffer or buffer-exchanged using Amicon Ultra-0.5 Centrifugal Filters with a 30 kDa MWCO.

### Native mass spectrometry

Native mass spectrometry measurements were performed using a hybrid quadrupole-time of flight mass spectrometer (Q-ToF Ultima, Micromass) equipped with a nano-electrospray ionization (nESI) source. Briefly, platinum-coated nESI emitters (internal diameter 1 to 3 μm) containing 3 to 5 μL of analyte solution were positioned approximately 3 mm from the heated (70 °C) sampling interface of the mass spectrometer^34^. Application of a 1.0 to 1.2 kV potential (positive mode) between the emitter tip and the sampling interface was used to generate gas-phase protein complexes. Instrument tuning settings and pressures were varied to minimize in-source dissociation of gas-phase complexes while maintaining sufficient ion transmission and detection^35^. Spectra were processed using vendor software (minimal smoothing using a Savitzky-Golay filter) and deconvoluted with UniDec^36^.

### Determination of protein switch *K*_*d*_ values using a FRET assay

The *K*_*d*_ values of the protein switches were measured using a FRET assay adapted from the protocol developed by Song *et al*.^25^ Mixtures of the CFP-E3-Switch and YFP-K3-Switch proteins were prepared, with the CFP-E3-Switch at a constant concentration of 0.1 μM and varying concentrations of K3-Switch-mVenus from 0.025 to 0.8 μM in a HEPES-based phosphorylation buffer (50 mM HEPES, 50 mM sodium chloride, 1 mM MgCl_2_, pH 7.5) supplemented with 100 μM ATP. Subsequently, 40 μL of each mixture were transferred to the wells of a 384-well flat bottom plate black with a transparent bottom in two series of wells. In the first series, 1 μL of water was added to each well. In the second series, 1 μL of recombinant PKA (1 mg/mL) was added to each well. In addition, two series of control wells were prepared: one series that contained only 0.025 to 0.8 μM of YFP-K3-Switch in phosphorylation buffer and another that contained only 0.1 μM CFP-E3-Switch. The plate was incubated for two hours at room temperature to complete the phosphorylation of protein switches mixed with PKA. The fluorescence emission spectra of all samples were recorded from 458 nm to 600 nm with 1 nm increments at the excitation wavelength of 433 nm (± 10 nm) on a CLARIOstar Plus plate reader (BMG LABTECH). The spectra were smoothed by using a 7-point moving average, and the background emission of CFP-E3-Switch (due to its emission at 533 nm) and of YFP-K3-Switch (due to its unspecific excitation at 433) nm in the control samples were subtracted from the corresponding samples containing the CFP-E3-Switch and YFP-K3-Switch mixtures to obtain the true FRET emission spectra. The FRET emission value at 533 nm was plotted as a function of the concentration of K3-Switch-mVenus. Non-linear regression analysis was performed in GraphPad Prism to calculate the *K*_*d*_ and 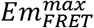 using Equation 8. The same procedure was followed to measure the *K*_*d*_ value of the E3_A_/K3-Switch, except the concentration of CFP-E3_A_-Switch was 0.5 μM and YFP-K3-Switch was varied from 0.1 to 5 μM.

### Real time FRET assays

For real time FRET measurements, CFP-E3_A_-Switch and YFP-K3-Switch were mixed in a 1:10 ratio (0.1 μM and 1 μM concentrations, respectively) in a HEPES-based phosphorylation buffer (50 mM HEPES, 50 mM sodium chloride, 1 mM MgCl_2_, pH 7.5) complemented with various concentrations of recombinant PKA as detailed in the main text. The phosphorylation reactions were started by adding 1 μL of 5 mM ATP to a 49 μL volume of the E3_A_/K3-Switch mixtures (100 μM final ATP) in the wells of a 384-well flat bottom black plate with a transparent bottom and in duplicates. Two fluorescence values were recorded in real time at the excitation value of 433 nm (± 10 nm) and the emission values of 475 nm (± 10 nm) corresponding to CFP emission and 533 nm (± 10 nm) corresponding to YFP and/or FRET emissions. The ratiometric FRET was calculated as the ratio of emissions at 533 nm and 475 nm and the mean value of each duplicate was plotted in GraphPad Prism. In experiments involving dephosphorylation, the phosphorylation buffer was complemented with 1 mM MnCl_2_ as λPP is dependent on manganese. For some experiments, the initial velocity *ν*_0_ of FRET production was calculated by non-linear regression in GraphPad Prism using equation 10.

## Declarations

The authors declare no competing interests.

## Acknowledgments

Protein production was performed at the Recombinant Products Facility (UNSW Sydney) and mass spectrometry experiments were performed at the Bioanalytical Mass Spectrometry Facility at the Mark Wainwright Analytical Centre (UNSW Sydney). This work was supported by the Air Force Office of Scientific Research (FA9550-17-1-0451). W.A.D. acknowledges support from the ARC (DP190103298 and FT200100798). D.L.W. was supported by the CSIRO Synthetic Biology Future Science Platform.

