## Supplemental data and figures for "Protein interaction kinetics delimit the performance of phosphorylation-driven protein switches"

Supplemental data for “**Protein interaction kinetics delimit the performance of phosphorylation-driven protein switches.**”

Daniel L. Winter, Adelgisa R. Wairara, Jack L. Bennett, William A. Donald, and Dominic J. Glover

Figure S1 ....... Predicted dynamic ranges at different concentrations of protein switches.

Figure S2 ....... Kinetic model of a two-component protein switch whose interaction is modulated by two phosphorylation sites.

Figure S3 ....... The reversibility of a phosphorylation-driven protein switch is limited by background kinase activity.

Figure S4 ....... Protein switch interaction kinetics rate modulate the negative feedback loop effect.

Figure S5 ....... E3/K3-Switch forms heterodimers as detected by native mass spectrometry.

Figure S6 ....... Simulated FRET response of $E3_A$/K3-Switch assuming dimerization as the sole mechanism of response to phosphorylation.

Figure S7 ....... Predicted response to phosphorylation of E3/K3-Switch and $E3_A$/K3-Switch based on the equilibria model.

Figure S8 ....... Ratiometric FRET (rFRET) response of E3/K3-Switch and of E3A/K3-Switch to PKA-mediated phosphorylation.

Figure S9 ....... Predicted increases in FRET upon phosphorylation of two-component protein switches at experimentally relevant concentrations.

Table S1 ........ Amino acid sequences of the phosphorylation driven protein switches

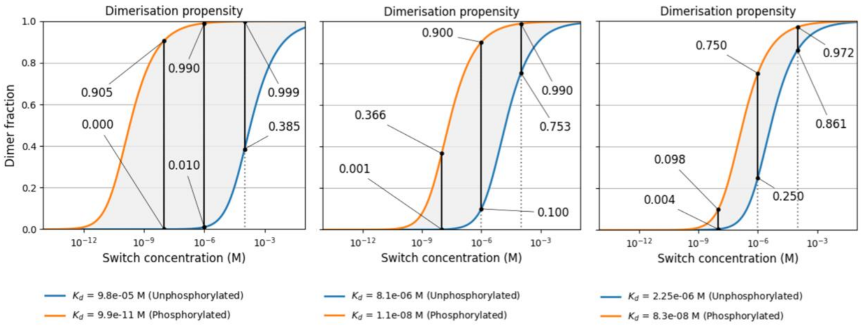


**Figure S1 – Predicted dynamic ranges at different concentrations of protein switches.** Three pairs of $K_d$ values corresponding to the unphosphorylated and phosphorylated states of protein switches that theoretically realize diverse dynamic ranges are represented. For each pair of $K_d$ values, the dimer fraction as a function of the protein switch concentration is plotted in blue for the theoretical unphosphorylated state and in orange for the theoretical phosphorylated state. The dynamic ranges at 10 nM, 1 µM and 100 µM protein switch concentrations are represented by vertical black bars, and the dimer fractions for both phosphorylation states are highlighted for each concentration.

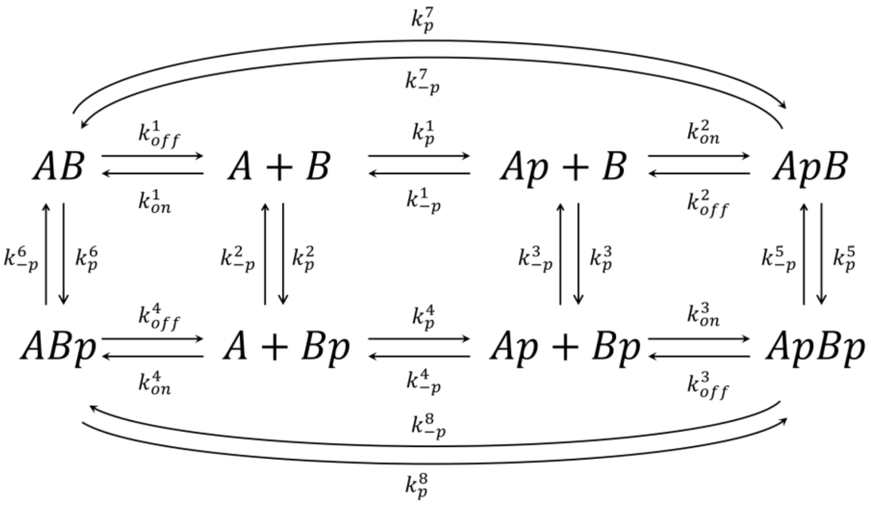


**Figure S2 – Kinetic model of a two-component protein switch whose interaction is modulated by two phosphorylation sites.** The dimerization of a two-component, heterodimeric protein switch (subunits denoted A and B) is governed by 24 kinetic parameters if it contains one phosphorylation site per subunit, and both phosphorylation sites affect dimerization; these consist of 8 binding kinetic parameters ($k_{on}$ and $k_{off}$) and 16 phosphorylation kinetic parameters ($k_p$ and $k_{-p}$). Note that phosphorylation events ($k_p$) are catalyzed by kinases require ATP as co-substrate and produce ADP as co-product, whereas dephosphorylation events ($k_{-p}$) are catalyzed by phosphatases and not require a co-substrate (but produce inorganic phosphate as co-product). Thus, phosphorylation and dephosphorylation are not the true reverse reactions of each other.

**Commented [DG1]:** I know it's a pain, but could the Kd values be updated for scientific notation? E.g. 9.9 x10-11.

Perhaps just replate/cover the text with the new notation

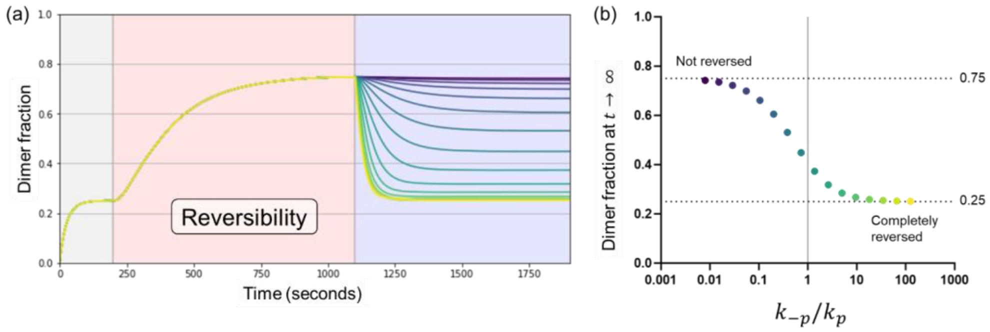


**Figure S3 – The reversibility of a phosphorylation-driven protein switch is limited by background kinase activity.** In a system where a protein switch is equilibrated (grey background), subjected to kinase activity (red background), and then to competing dephosphorylation activity (blue background), phosphatase activity must vastly exceed kinase activity to completely revert the protein switch to its baseline activity. **(a)** Kinetic model simulations where the phosphatase activity is varied from 125 times lower (purple graph) to 125 higher (yellow graph) than the competing kinase activity. The simulations are the same as those depicted in main Figure 4b except that additional $k_p^{dimer}$ and $k_{-p}^{dimer}$values are plotted for a total of 16 graphs. **(b)** Plot of the dimer fractions at $t \to \infty$ as a function of the ratio between phosphatase activity ($k_{-p}$) and kinase activity ($k_p$). The monomer dimer baselines of the protein switch (0.25 and 0.75, respectively) are indicated as dotted lines. The colors of the data points correspond to the simulation graphs in (a). In these simulations, it is assumed that both the kinase and the phosphatase have equal activities towards the monomeric and dimeric forms of the protein switch and that ATP cannot be depleted.

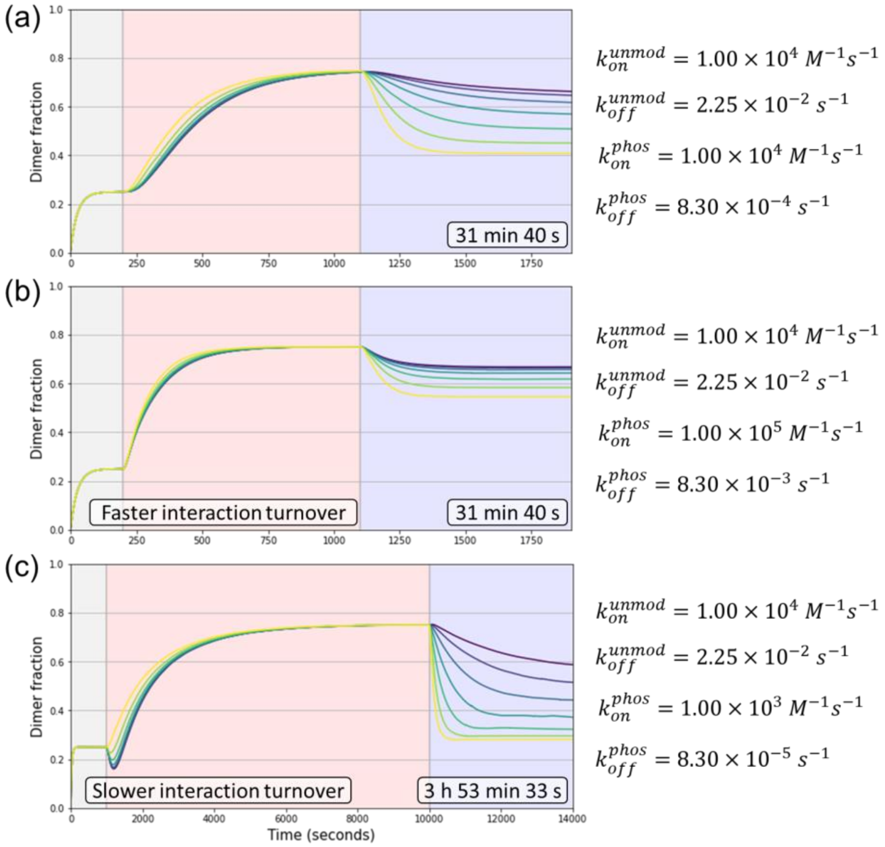


**Figure S4 – Protein switch interaction kinetics rate modulate the negative feedback loop effect.** The predicted phosphorylation-driven dimerization of a two-component protein switch are shown for varying $k_{on}^{phos}$ and $k_{off}^{phos}$ rates that result in the same dissociation constant of and $K_d^{phos} = 8.30 \times 10^{-8}\ M$ after phosphorylation. In all simulations, $k_{on}^{unmod}$ and $k_{off}^{unmod}$ are constant and the phosphorylation kinetics of monomeric form of the protein switch are constant as well ($k_p^{monomer} = k_{-p}^{monomer} = 1 \times 10^{-2} s^{-1}$). The phosphorylation kinetics of the dimeric form ($k_p^{dimer}$ and $k_{-p}^{dimer}$) of the protein switch are varied as indicated in Figure 4c of the main document, illustrating situations where dimerization does not affect (yellow graphs) or greatly attenuates the likelihood of a switch to be (de)phosphorylated 100-fold (purple graphs). **(a)** The data from Figure 4c is shown as reference, which illustrates the "negative feedback" effect whereby dimerization of the protein switch attenuates reversibility when the phosphorylation site is occluded in the switch's dimeric form. The additional simulations show that **(b)** faster interaction turnover rates (higher $k_{off}^{phos}$ and $k_{on}^{phos}$) further exacerbate the negative feedback. Additionally, the response time of the switch is improved; note that the equilibrium is reached substantially faster in the phosphorylation stage of the simulation when compared to (a). In contrast, **(c)** slower interaction turnover rates (lower $k_{off}^{phos}$ and $k_{on}^{phos}$) improve the reversibility of the switch, but greatly affect the response time to phosphorylation; note that the simulated time spans nearly 4 hours of protein interaction dynamics, instead of approximately 30 minutes as in (a) and (b). Moreover, the switch responds to phosphorylation with an initial decrease in the dimer fraction before dimerization increases. The graph background colors indicate the equilibration (grey), phosphorylation (red), and dephosphorylation (blue) stages of the simulation; during the third stage of the simulation, the kinase activity persists and competes with phosphatase activity.

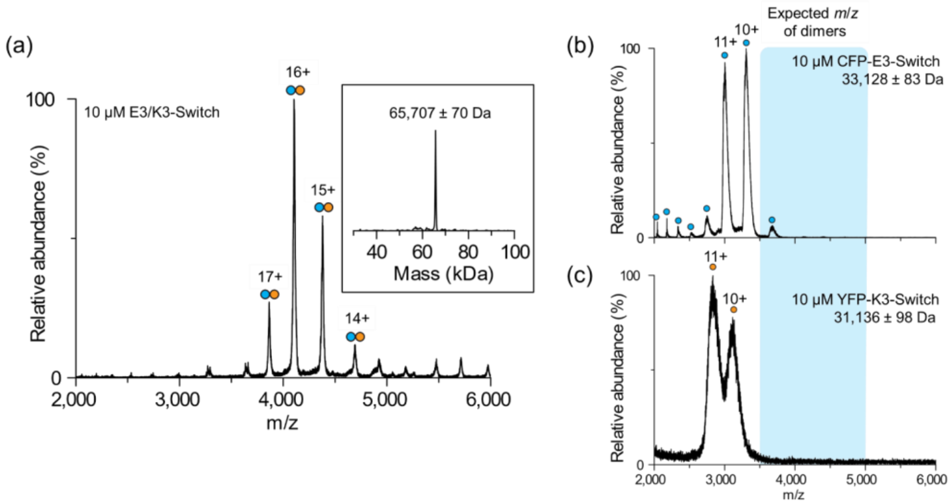


**Figure S5 – E3/K3-Switch forms heterodimers as detected by native mass spectrometry. (a)** Native mass spectrum of the E3/K3-Switch heterodimer (10 µM each subunit, represented as blue and orange disks) in 200 mM ammonium acetate, pH 7. The inset, which displays the deconvoluted (zero-charge) mass spectrum, shows the predominant formation of dimers. **(b, c)** Neither CFP-E3-Switch nor YFP-K3-Switch forms homodimers at experimentally relevant concentrations. Native mass spectra of (b) 10 µM CFP-E3-Switch (blue disks), and (c) 10 µM YFP-K3-Switch (orange disks) in 200 mM ammonium acetate, pH 7. Signals corresponding to homodimeric species could not be observed for either protein switch subunit.

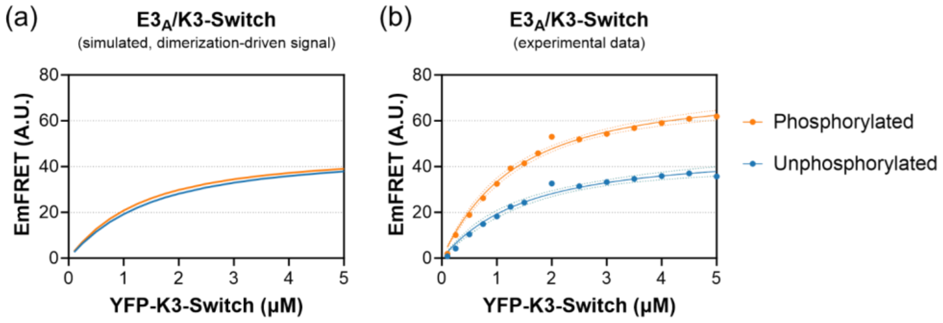


**Figure S6 – Simulated FRET response of E3A/K3-Switch assuming dimerization as the sole mechanism of response to phosphorylation. (a)** Simulated FRET response of $E3_A$/K3-Switch assuming a concentration of 0.5 µM CFP-$E3_A$-Switch and the experimentally measured $K_d^{unmod} = 1.171\ \mu M$ and $K_d^{phos} = 0.9989\ \mu M$ values from experimental data and an $Em_{FRET}^{max} = 47.53$ value measured from experimental data for the unphosphorylated protein switch. **(b)** The experimental data (blue and orange points), also shown in figure 5b of the main text, for comparison. The $Em_{FRET}^{max}$ value for the phosphorylated state of the $E3_A$/K3-Switch, calculated by non-linear regression (blue and orange lines), is much higher (see Table 3 in the main text for all best fitted parameters). The experiment reveals that mechanisms other than dimerization drive the FRET response of the $E3_A$/K3-Switch.

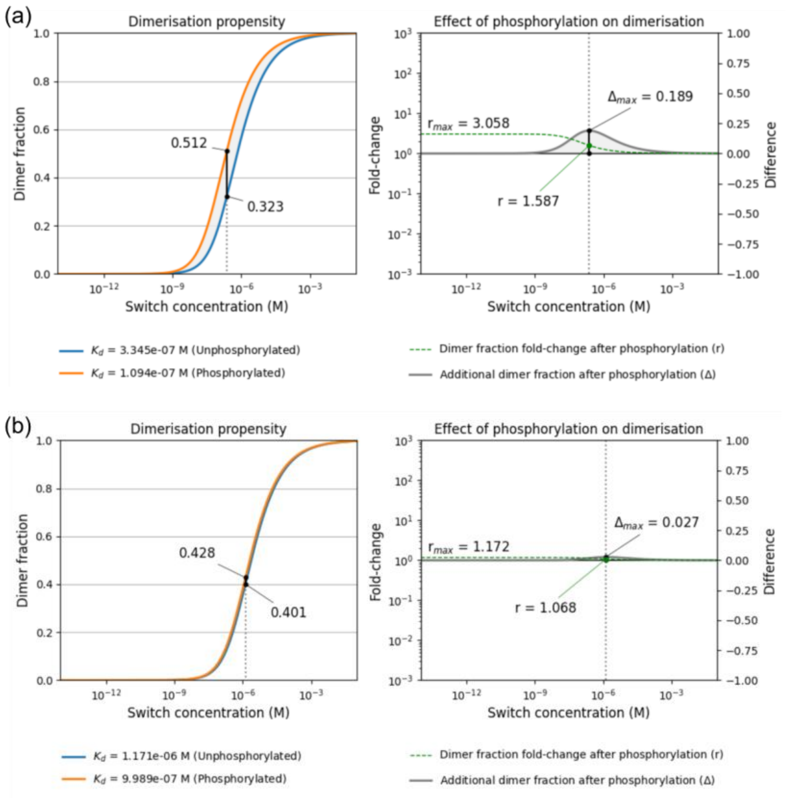


**Figure S7 – Predicted response to phosphorylation of E3/K3-Switch and $E3_A$/K3-Switch based on the equilibria model. (a)** On the left, the calculated dimeric fraction as function of the E3/K3-Switch in its unphosphorylated state (blue graph) and completely phosphorylate state (orange graph). On the right, the calculated difference (Δ, grey graph) and fold-change ($r$, green, dotted graph) in dimer fraction in response to complete phosphorylation and as a function of concentration). **(b)** Same as in (a) but for the $E3_A$/K3-Switch. The black vertical lines in each panel indicate at what concentration the predicted greatest absolute change in dimer concentration $\Delta_{max}$ occurs for each switch.

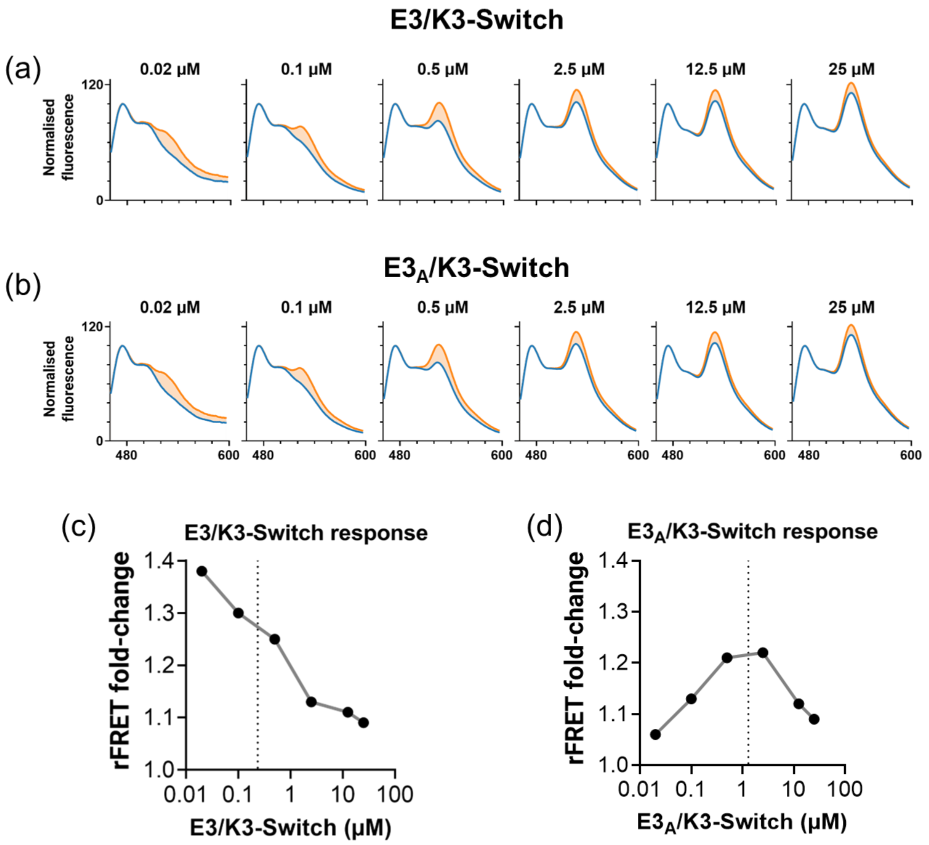


**Figure S8 – Ratiometric FRET (rFRET) response of *E3/K3-Switch and of E3$_A$/K3-Switch to PKA-mediated* phosphorylation. (a)** FRET response of E3/K3-Switch and **(b)** of E3$_A$/K3-Switch at different switch concentrations. FRET emission spectra in the non-phosphorylated state are shown in blue, and FRET emission spectra in the phosphorylated state are shown in orange, from 460 nm to 600 nm emission wavelengths. The filled area between each pair of graphs (light orange) corresponds to the additional FRET signal resulting from phosphorylation-driven dimerization of the protein switch. **(c, d)** Fold-change in rFRET upon phosphorylation of each protein switch calculated for each concentration from the data in (a, b). The dotted line indicates the predicted concentration for the greatest change in dimer fraction ($D^{phos} - D^{unmod}$, see Figure S7).

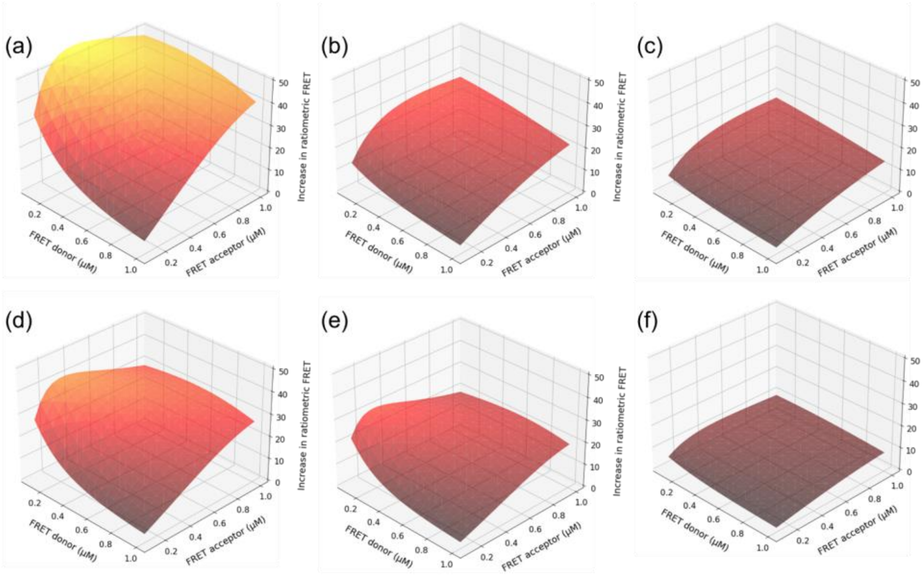


**Figure S8 – Predicted increases in FRET upon phosphorylation of two-component protein switches at experimentally relevant concentrations.** The increase in ratiometric FRET signal upon phosphorylation of a protein switch is proportional to the increase in the fraction of the FRET donor subunit (e.g., CFP-E3-Switch) bound to the FRET acceptor (e.g., YFP-K3-Switch). The surface plots show the theoretical increase in FRET signal for different pairs of $K_d^{unmod}$ and $K_d^{phos}$ values at experimentally relevant concentrations of the two subunits, with each subunit varying in concentration from 0.1 μM to 1 μM. The surface plot is colored following a heatmap (from dark red to yellow) to easily visualize the height of the surface along the Z axis representing the increase in FRET signal. Note that the Z maxima are often found for X and Y values where the concentration of the FRET acceptor is higher than the FRET donor. Each plot represents the predicted FRET response to phosphorylation for a two-component protein switch whose $K_d$ value decreases in response to phosphorylation **(a)** from $K_d$ = 1 μM to 0.1 μM; **(b)** from $K_d$ = 1 μM to 0.333 μM; **(c)** from $K_d$ = 1 μM to 0.5 μM; **(d)** from $K_d$ = 0.5 μM to 0.1 μM; **(e)** from $K_d$ = 0.333 μM to 0.1 μM; and **(f)** from $K_d$ = 05 μM to 0.333 μM.

**Table S1 - Amino acid sequences of the phosphorylation driven protein switches[a]**

| | |
|---|---|
| CFP-E3-Switch | MVSKGEELFTGVVPILVELDGDVNGHKFSVSGEGEGDATYGKLTLKFICTTGKLPVPWPTLV<br>TTLSWGVQCFARYPDHMKQHDFFKSAMPEGYVQERTIFFKDDGNYKTRAEVKFEGDTLVNRI<br>ELKGIDFKEDGNILGHKLEYNAIHGNVYITADKQKNGIKANFGLNCNIEDGSVQLADHYQQN<br>TPIGDGPVLLPDNHYLSTQSKLSKDPNEKRDHMVLLEFVTAAGITLGMDELYSGENLYFQGG<br>*gabcdefgabcdefgabcdefgabcdefgabcdef*<br>SG**EIAALEKEIAALEKRARRGSARVRELENEIAALEK**GGSGHHHHHH |
| CFP-E3$_A$-Switch | MVSKGEELFTGVVPILVELDGDVNGHKFSVSGEGEGDATYGKLTLKFICTTGKLPVPWPTLV<br>TTLSWGVQCFARYPDHMKQHDFFKSAMPEGYVQERTIFFKDDGNYKTRAEVKFEGDTLVNRI<br>ELKGIDFKEDGNILGHKLEYNAIHGNVYITADKQKNGIKANFGLNCNIEDGSVQLADHYQQN<br>TPIGDGPVLLPDNHYLSTQSKLSKDPNEKRDHMVLLEFVTAAGITLGMDELYSGENLYFQGG<br>*gabcdefgabcdefgabcdefgabcdefgabcdef*<br>SG**EIAAAEKEIAALEKRARRGSARVRELENEIAALEK**GGSGHHHHHH |
| YFP-K3-Switch | MVSKGEELFTGVVPILVELDGDVNGHKFSVSGEGEGDATYGKLTLKLICTTGKLPVPWPTLV<br>TTLGYGLQCFARYPDHMKQHDFFKSAMPEGYVQERTIFFKDDGNYKTRAEVKFEGDTLVNRI<br>ELKGIDFKEDGNILGHKLEYNYNSHNVYITADKQKNGIKANFKIRHNIEDGGVQLADHYQQN<br>TPIGDGPVLLPDNHYLSYQSKLSKDPNEKRDHMVLLEFVTAAGITLGMDELYSGENLYFQGG<br>*gabcdefgabcdefgabcdefgabcdefgabcdef*<br>SG**KIAALKEKIAALKERARRGSARVRELENKIAALKE**GGSGHHHHHH |

[a]The kinase-sensitive coiled coil sensors are shown in bold, underline font; the protein kinase A target motif is highlighted in yellow; the coiled coil register (location of residues within the coiled coil heptad) is shown above the coiled coil sequences in lower case.
